# Combinatorial organoid mutagenesis screen reveals gene constellations driving malignant transformation, pathology and chemosensitivity in high-grade serous ovarian carcinoma

**DOI:** 10.1101/2025.03.10.642422

**Authors:** Daryl J. Phuong, Coulter Q. Ralston, Tony M. Ezzat, Christopher S. Ashe, Amanda P. Armstrong, Andrea Flesken-Nikitin, Robert J. Yamulla, Alexander Yu. Nikitin, John C. Schimenti

## Abstract

High-grade serous ovarian carcinoma (HGSC) is the sixth leading cause of cancer-related death among women. Many cases arise from the Fallopian tubal epithelium (TE), exhibit numerous mutations, and present heterogenous pathological features. However, the contribution of specific mutation combinations to cellular transformation, pathological phenotype and chemotherapeutic response remains poorly understood. Here, we used a *Trp53*-deficient mouse TE-derived organoid platform to perform combinatorial CRISPR mutagenesis of 20 candidate HGSC driver genes. Mutations in *Nf1*, *Cdkn2a* and *Map2k4* were most frequently observed in transformed organoids. Upon transplantation into mice, those containing *Map2k4* mutations predominantly gave rise to papillary-glandular histology, whereas those containing *Nf1* mutations formed more mesenchymal-like carcinomas. Transcriptomic analysis revealed that *Nf1*-mutant tumors of all pathological phenotypes overexpressed the long non-coding RNA *Pvt1*, a marker associated with poor prognosis in HGSC patients. *Map2k4*-mutant organoids were more sensitive to paclitaxel and niraparib, while *Nf1*-mutant combinations responded better to trametinib. Notably, the removal of Rho kinase inhibitor (ROCKi) reduced trametinib sensitivity in both *Map2k4*– and *Nf1*-mutant organoids, underscoring the importance of culture conditions and potential antagonistic drug interactions in organoid-based drug screens. Collectively, our results demonstrate that TE-derived organoids coupled with combinatorial CRISPR mutagenesis provide a powerful system to unravel the genetic and phenotypic complexity of HGSC. In particular, we found that *Map2k4* functions as a tumor suppressor that shapes distinct tumor histology and chemosensitivity, suggesting it as a potential therapeutic target in select HGSC cases.

## INTRODUCTION

The transformation of normal cells into cancer cells is driven by multiple genetic events. However, pinpointing the exact combination of mutations that fuel cancer progression and influence treatment responses is challenging. High-grade serous carcinoma (HGSC) accounts for over 70% of ovarian cancer deaths (1), and TCGA data (available at cBioPortal.org) reveal an average of 46 mutations per HGSC sample. Chromosomal rearrangements that disrupt key tumor suppressors like *RB1, NF1, RAD51B*, and *PTEN* have been valuable for identifying key drivers of HGSC (1–8). Furthermore, combinations of somatic and germline mutations play a critical role in determining patient outcomes, tumor characteristics, and immune responses, highlighting why sequencing individual tumors is key to personalized treatment (1,5). Although *TP53* mutations are present in >90% of HGSCs, *Trp53* ablation alone is insufficient for HGSC induction in mice; alterations of 1-3 additional genes are necessary (8–11).

HGSC originates from both ovarian surface epithelium (OSE) and tubal epithelium (TE) cells (7,12–15), but whether they share the same genetic drivers for malignant transformation is unclear. Studies show up to 60% of HGSCs are linked to serous tubal intraepithelial carcinoma (STIC) lesions (16–20), with STICs found in 8% of patients with *BRCA1/2* mutations undergoing risk-reducing salpingo-oophorectomy (18,21–25). Although HGSC was originally believed to only be of OSE origin, mounting evidence suggests that the preponderance of cases arise from the TE (2,20,26).

Organoids are a more biologically relevant model than traditional 2D cell cultures for studying cancer biology. Mouse OSE and TE organoids have been used to identify HGSC cell origins and to validate key genetic drivers like *Trp53* and *Rb1* (10,11,27). For example, TE organoids with *Trp53*, *Brca1*, and *Nf1* mutations showed higher tumorigenic potential than OSE organoids with the same mutations (10), supporting the notion that tumors arising from different cell types are driven by distinct mutational landscapes and cell interactions (3,28). The complexity of mutational load in different HGSC cases challenges the practical use of predefined gene combinations, especially in genetically engineered mouse models that acquire secondary mutations similar to those in humans (29). Organoids offer a platform to explore additional genes, mutation combinations, and drug sensitivities in HGSC.

HGSC is classified into four molecular subtypes associated with tumor progression – immunoreactive, proliferative, mesenchymal, and differentiated – though their prognostic value for patient survival remains unclear (30–35). Pathologically, HGSC can present in various patterns the most common of which are papillary, glandular, solid, or SET (solid, pseudoendometrioid, transitional) (31,36,37). *Brca1* mutations are often associated with SET tumors and the immunoreactive subtype (38–40). However, over 70% of HGSC cases are diagnosed at advanced stages, complicating the identification of early genetic drivers and their impact on subtypes and histology (41–43).

To identify genetic drivers of TE transformation, we coupled an organoid platform with random combinatorial mutagenesis. This approach allowed us to induce mutations in seven or more HGSC-associated genes simultaneously, and to identify combinations that conferred sensitivity or resistance to transformation. Mutations in *Nf1*, *Cdkn2a* and *Map2k4* were the most common in transformed organoids, and we found distinct organoid growth patterns, chemoresistance features, tumor pathologies and transcriptome profiles characteristic of *Nf1* and *Map2k4* deficiencies.

## RESULTS

### Approach for screening candidate HGSC suppressors in a TE organoid model

We leveraged a combinatorial CRISPR/Cas9 mutagenesis screen in TE organoids as a high-throughput platform for identifying mutation combinations that drive HGSC. TE organoids offer several important properties for this application: they can form from single cells (Sup. Fig. 1A, B), express key differentiation markers (acetylated alpha-tubulin/FOXJ1 for ciliated cells, OVGP1 for secretory cells, and PAX8 for immature cells) (Sup. Fig. 1C) (10,11,15,44), and if genetically transformed, they exhibit morphological changes linked to increased tumorigenic potential in mice (10,11,44). The mutagenesis screen involved infecting individual TE organoid-derived cells (derived from the uterine tube, also known as oviduct in mice and fallopian tube in humans) with a lentiviral library at high multiplicity of infection (MOI), then allowing them to form new organoids. Those with abnormal morphologies were selected, and the gRNAs borne by each were detected (Fig. 1). Prior to screening, the gRNA distribution of the library was determined as a baseline for identification of enriched targeting vectors in aberrant organoid clones (Sup. Fig. 1D, E). We confirmed that secondary organoids were clonal, since cells independently infected with GFP or mCherry reporters formed organoids without dual reporter expression when combined in culture (Sup. Fig. 1B).

**Figure 1.**
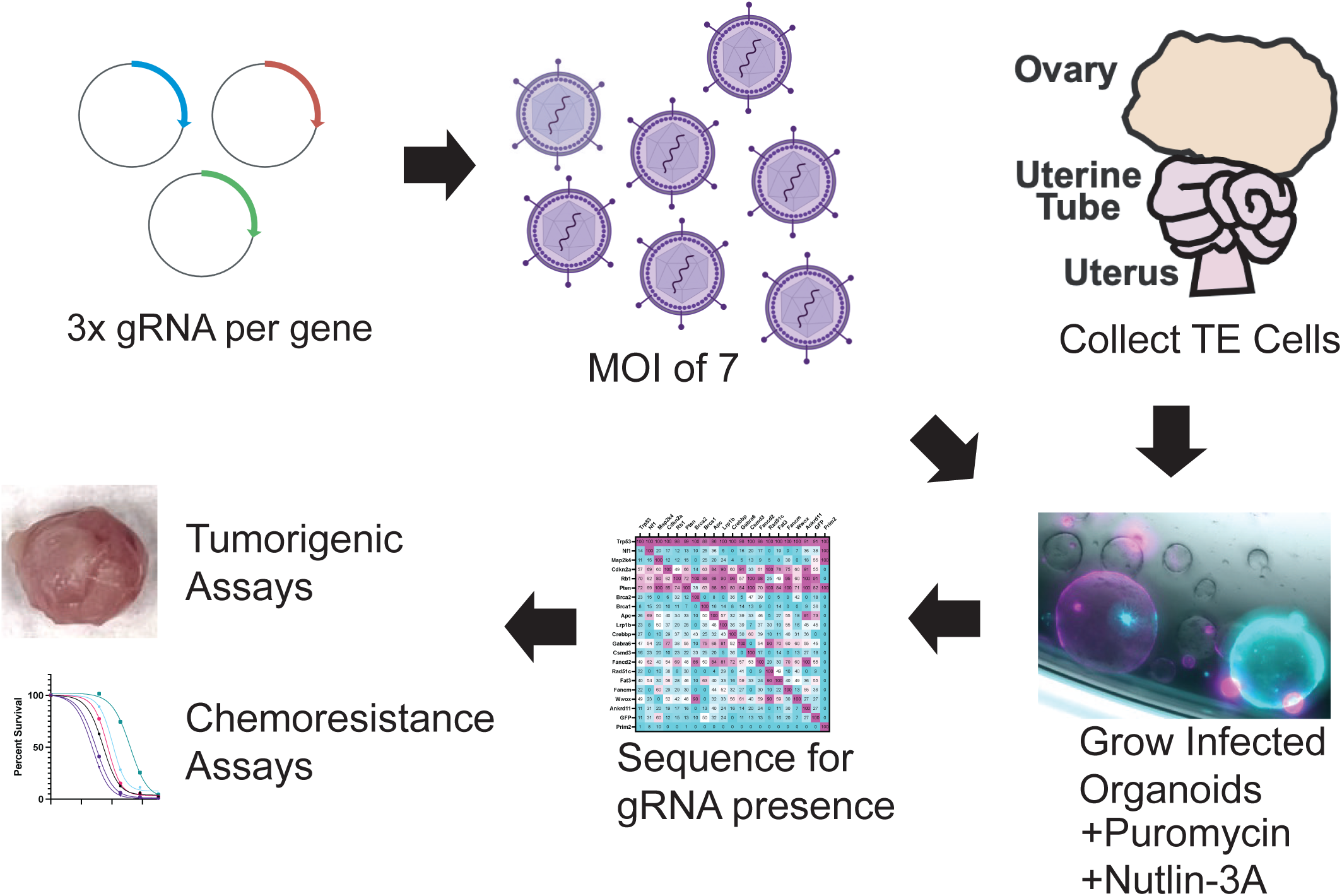
Schematic of screening strategy. Tubal epithelial cells (TE) are from the uterine tube.

### Identification of target genes enriched in aberrant organoids

We designed the screen to target 20 genes, 18 of which are commonly mutated in HGSC (based on TCGA data) and two others, *Fancm* and *Apc*, that are also cancer-related (33,45). A lentiviral library consisted of three gRNAs per gene was cloned into a plasmid vector (lentiCRISPRv2) designed to express both the gRNA and Cas9, packaged as lentiviruses, and used to infect TE organoid-forming cells at an estimated multiplicity of infection (MOI) of 7 (Fig. 1). Given that *TP53* mutations are found in over 90% of HGSC cases, the organoids were grown in Nutlin-3a to select for *Trp53*-deficient cells (Fig. 1) (45). After 14 days, we visually screened for abnormal organoids – those with folds, grapelike structures, protrusions, solid formations, or larger sizes compared to controls lacking only *Trp53* (Fig. 2A). The fraction of aberrant organoids was 0.28% ± SD = 0.0311 (n=5 independent library infections). We then picked and expanded these organoids, extracted DNA to sequence vector amplicons, and found that individual organoids had an average of 8 gRNAs (Fig. 2B). Although we expected that all the clones would contain a gRNA targeting *Trp53* due to Nutlin-3A selection, 11 of 95 did not; it is possible that the lentivirus did not stably integrate in these clones (yet they survived short term puromycin selection), or that *Trp53* was poorly expressed or inactivated by other means in these organoids.

**Figure 2.**
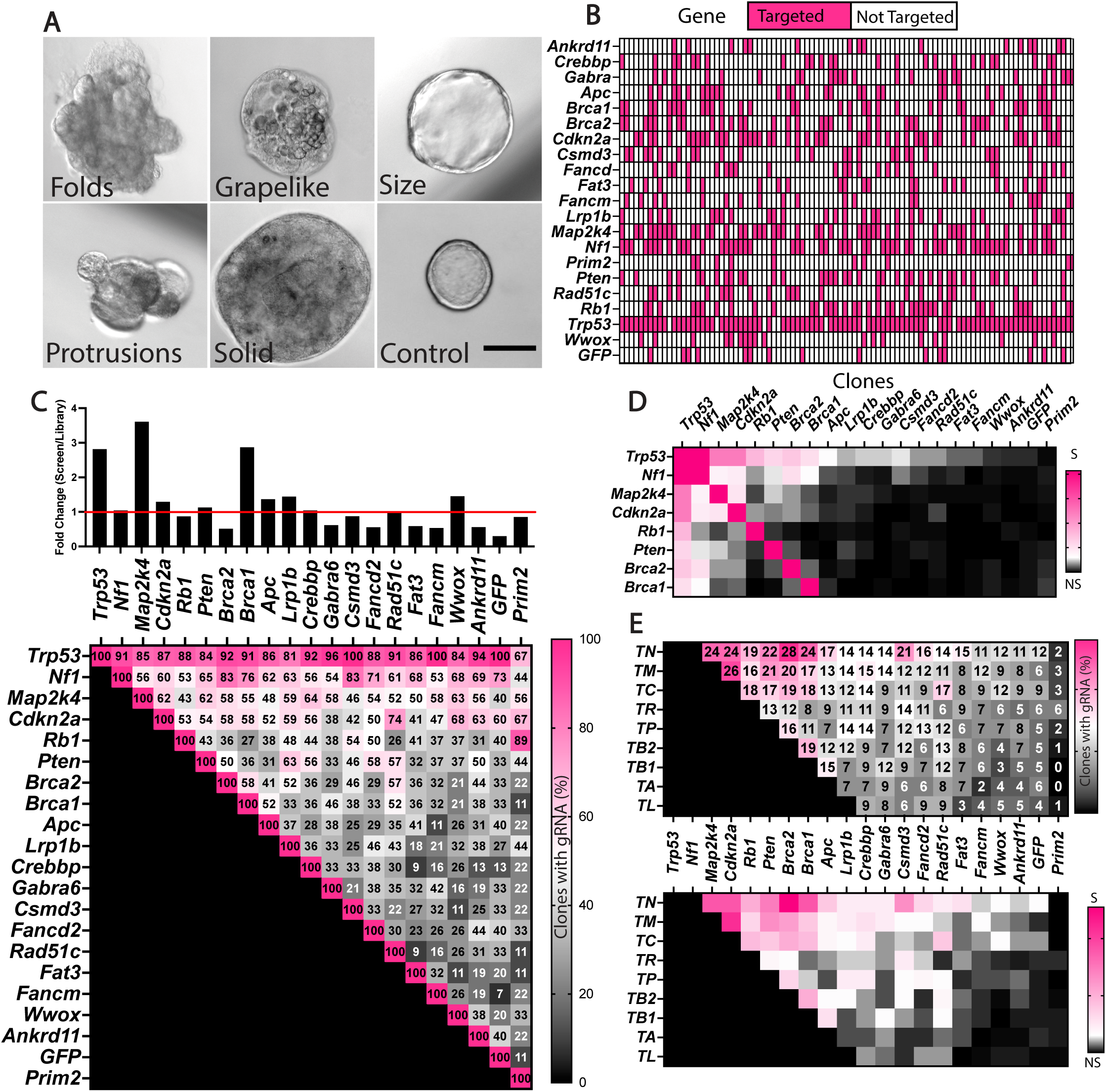
Mutation combinations in transformed TE organoids. A) Phenotypes of mutant organoids. Scale bar = 100µm. B) gRNA vectors identified in each aberrant organoid clone. C) Enrichment of targeting vectors in aberrant clones. The fold change was calculated based on % representation in clonal organoids over % representation in library. The red horizontal line represents no representation difference between gRNA frequency found in library and screen. (Bottom) Frequency of co-mutations in organoids. Numbers are percentages. D) ξ2 analysis of significant TE double mutations, DF, 19, P<0.05%, S= Significant, NS= not significant. E) Number of clones that contain gRNA’s targeting genes in triple mutations containing the top 9 overrepresented genes in the screen. (Bottom) ξ2 analysis of significant TE triple mutations, DF, 19, P<0.05%. S= Significant, NS= Not Significant. *Trp53*, abbreviated (“T”), *Nf1* (N), *Map2k4* (M), *Cdkn2a* (C), *Rb1* (R), *Pten* (P), *Brca2* (B2) and *Brca1* (B1).

After adjusting for vector representation in the plasmid library used to package lentivirus, the most dramatically enriched gRNAs corresponded to *Map2k4* and *Brca1* (Fig. 2C), whereas several were highly underrepresented, notably *Brca2*. The underrepresentation of other vectors may reflect either a compromise to viability or a dependency of co-mutation with certain other genes in order to confer abnormal organoid phenotypes. For example, *Nf1* and *Cdkn2a*, which were not enriched when considering their overall normalized representation (Fig. 2C), yet they were amongst the most statistically enriched in pairwise combinations with additional mutations (Fig. 2D). The most enriched triple mutant combinations (all including *Trp53,* abbreviated “T”) involved *Nf1* (N)*, Map2k4* (M), *Cdkn2a* (C)*, Rb1* (R)*, Pten* (P)*, Brca2* (B2) and *Brca1* (B1) and will henceforth be abbreviated as indicated when in combinations (Fig. 2D, E).

### Key mutation combinations driving TE transformation

To validate the combinatorial contribution of gene disruptions to transformation, we infected organoid-forming TE cells with specific pairs or trios of lentiviral vectors. Trp53/*Nf1* or Trp53/*Pten* co-mutations the fraction of aberrant organoids over p53 alone (Fig. 3A, B), consistent with previous findings that loss of *Nf1* and *Pten* drives HGSC (44). Trio mutations including *Nf1* resulted in larger organoids with intact lumens, while those including *Pten* caused folds with blob-like protrusions and denser cores (Fig. 3E). In contrast, other double or triple mutant combinations (such as some involving *Brca1/2*) reduced organoid diameter and formation rate (Fig. 3A-D), consistent with previous data suggesting homozygous *Brca1* loss is rare in rapidly growing organoid systems (10).

**Figure 3.**
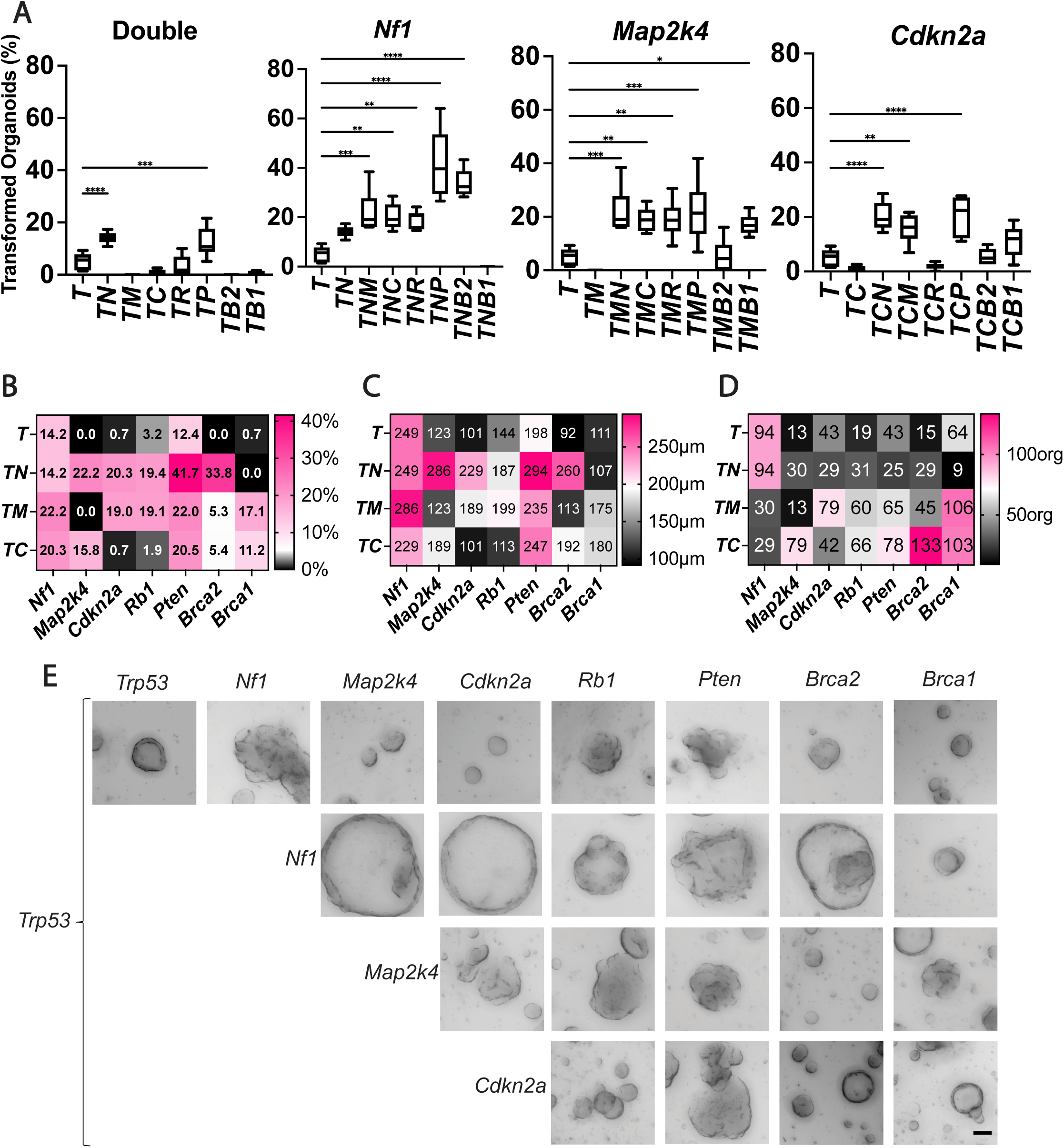
Validation of mutation combinations driving TE organoid transformation. Percent transformation of A) Double mutant combinations, Triple mutant combinations with Nf1^mut^, Map2k4^mut^ or Cdkn2a ^mut^ background. B) Organoid transformation percentage of combinations compared to Trp53^-/-^ alone. C) Average organoid diameters vs Trp53^-/-^ alone D) Average organoid (org) formation vs Trp53^-/-^ alone. E) Representative images of organoids for each combination tested. Scale bar = 100µm. One-way anova was used to assess significance, n=6 per condition. P-value * ≤ 0.05, ** ≤0.01, *** ≤0.001, **** ≤ 0.0001. *Trp53*, T; *Nf1*, N; *Map2k4,* M; *Cdkn2a,* C; *Rb1,* R; *Pten,* P; *Brca2,* B2; *Brca1,* B1.

*Trp53* mutations with homozygous *Nf1* and *Pten* deletions are present in 1.26% of human HGSCs (Firehose legacy data in cBioPortal), and TE organoids with these mutations form HGSC-like tumors upon transplantation (44). We generated specific triple mutant organoid combinations to validate the bulk (Fig. 2E) screen results. Disruption of genes besides *Pten* enhanced transformation with TN deficiency, particularly mutations in *Map2k4*, *Cdkn2a*, *Rb1*, or *Brca2*, which increased transformation compared to *Trp53* alone. However, only *Pten* or *Brca2* mutation markedly enhanced transformation in TN*-*deficient organoids (Fig. 3A, B). The TNB1 combination resulted in poor growth *ex vivo* (10) (Fig. 3C). Our data (Fig. 3A, B) supports previous findings showing that TNP disruption increased aberrant organoid formation rate (44). Interestingly, the phenotype of these organoids was a hybrid of those with TN and TP mutation combinations, exhibiting less overall folding but more ruffled edges and smaller blob-like protrusions (Fig. 3E).

The most overrepresented targeted gene in the screen was *Map2k4*, which acts in the JNK pathway and is considered a potential HGSC tumor suppressor (46,47). Despite this, *Map2k4* deletion remains to be functionally studied in mice or organoids for its involvement in ovarian cancer. According to data from the TCGA, *MAP2K4* mutations and homozygous deletions are found in 0.32% and 4.2% of HGSC cases, respectively (48). Consistent with these reports, *Map2k4* mutations increased transformation in the presence of other mutations except for *Brca2* in our screen. Although *Map2k4* mutations increased transformation, they had notably different effects than other mutation combinations. In fact, *Map2k4* mutations had less of an impact on organoid diameter than *Nf1* mutations (Fig. 3C, E). Furthermore, in triple mutation combinations, TMB1 mutants showed increased transformation, while TMB2 did not – opposite to the patterns seen in TNB1 and TNB2 (Fig. 3A, B). This aligns with the screen’s overrepresentation of *Map2k4* and *Brca1* (Fig. 2C).

We next investigated the role of *Cdkn2a*, as it was the third most co-mutated gene in triple mutation combinations with *Trp53* within the screen (Fig. 3E). *Cdkn2a* encodes p14 and p16 proteins, which are involved in the RB pathway and often used to distinguish HGSC from other ovarian cancers (49). Co-mutating *Cdkn2a* with *Trp53* reduced transformation potential, organoid size, and formation rate compared to *Trp53* alone (Fig. 3A-D). Adding *Rb1* to TC only slightly improved these parameters, while *Brca1/2* additions restored them to levels similar to *Trp53* alone. Collectively, the only *Nf1*, *Map2k4*, or *Pten* co-mutations further improved transformation (Fig. 3A-D).

The most significantly overrepresented triple mutation combinations were *Map2k4/Cdkn2a* and *Nf1/Pten* combined with *Trp53* (Sup. Fig. 4A, B). For transformed organoids, additional overrepresented mutations included [*Nf1, Pten, Brca2, Crebbp, Rad51, Wwox*] with TMC and [*Map2k4, Cdkn2a, Brca1, Lrp1b, Crebbp*] with TNP (Sup. Fig. 4A, B). Quadruple mutant combinations TNP*Rad51C* and TMCP enhanced transformation (Sup. Fig. 4C, D). Organoid formation frequency increased with co-mutation of *Map2k4, Cdkn2a, Brca1, Lrp1b*, or *Csmd3* in a TNP deficient background, and *Rb1* or *Csmd3* increased formation with TMC (Sup. Fig. 4C, D). Our findings suggest that while a fourth mutation does not significantly enhance transformation in most cases, it does not rule out roles in cell growth, survival, metastasis, and chemoresistance. Our goal was to identify the minimal genes required for aberrant organoid transformation, warranting further exploration of novel triple combinations.

### Mutation combinations drive differences in tumorigenic potential, gene expression and tumor pathology

To assess whether mutation combinations linked to aberrant organoid formation and morphology reflect *in vivo* tumor formation, we performed subcutaneous flank injections of organoid derived cells into immunodeficient (NSG) female mice and monitored them for up to 200 days. Loss of *Trp53* alone did not result in tumor formation, consistent with prior studies (2,5,7). Among the combinations tested, TNP led to the shortest survival, with tumors reaching 1 cm within an average of 70 days post-injection (Sup. Fig. 5A). In contrast, survival times were longer for TN (138 days) and TP (172 days) combinations (Sup. Fig. 5A). Adding *Brca2* (136 days) or *Map2k4* (124 days) to the TN background had minimal impact on survival (Sup. Fig. 5A).

In the context of TP deficiency, additional mutations in *Map2k4* or *Cdkn2a* reduced survival by 46 and 28 days, respectively, but tumor size doubling times increased to 62 and 48 days compared to 27 days for TP alone (Sup. Fig. 5A). Tumor latency mirrored these findings, with TPM and TPC averaging 96 and 95 days, respectively, compared to 138 days for TP (though this lacked statistical significance due to small sample size; Sup. Fig. 5B). Notably, organoids with TM co-mutations and additional losses in *Cdkn2a, Rb1,* or *Brca1* failed to efficiently form tumors within 200 days. Collectively, these results suggest that *Trp53* loss combined with either *Nf1* or *Pten* constitutes a minimal requirement for tumor formation, and additional losses of *Map2k4, Cdkn2a*, or *Brca2* enhance tumor growth and formation rates.

We next assessed whether organoid metrics – formation, transformation, and diameter (Fig. 3B– D) – predicted mouse survival after tumor induction via organoid-derived cell injections. In general, larger organoid diameter and faster transformation rate, but not formation efficiency, correlated with reduced survival (Sup. Fig. 5A, C-E). Only the TNP mutations had a significantly higher tumor formation rate and growth than TN or TP, leaving open the possibility that certain mutation combinations might impact tumor presentation.

We then assessed whether *Nf1* or *Map2k4* mutations would alter growth potential and survivability of mice. We found that *Nf1* mutations resulted in a faster time from initial tumor formation and reduced survivability compared to their *Map2k4* counterparts (Fig. 4AB). These findings were consistent with low *Nf1* expression in human samples resulting in reduced survivability (Sup. Fig. 5F).

**Figure 4.**
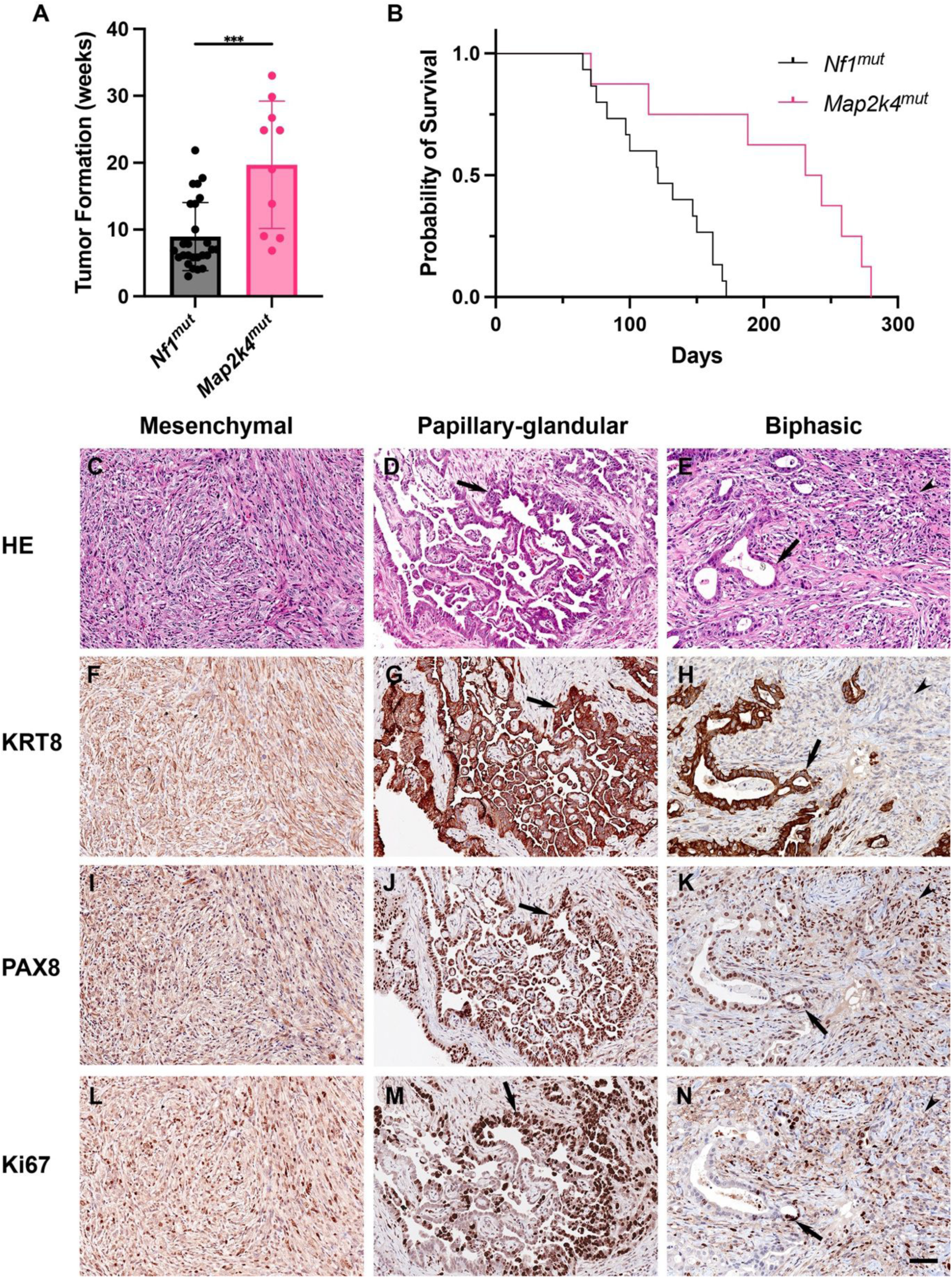
Characterization of neoplasms arising from genetically altered TE organoid cells transplanted subcutaneously. Weeks post injection until A) Tumor initiation and B) Mouse was sacrificed C) Mesenchymal (Mesenchymal-like) carcinoma with monotonous tight spindle-cell growth pattern, TN. D) Papillary-glandular carcinoma with slit-like spaces forming complex labyrinthine pattern (arrow), TMP. E) Biphasic carcinoma with both spindle-cell (arrowhead) and glandular (arrow) patterns of growth, TP. Hematoxylin and eosin (HE; C-E). F-N) Expression (brown color) of KRT8 (F-H), PAX8 (I-K), and Ki67 (L-N) in neoplastic cells. Counterstaining with hematoxylin (F-N). Scale bar, all panels 60 μm.

After subcutaneous transplantation, neoplasms arising from genetically altered TE organoids could be stratified into 3 groups (Fig. 4C-N., Sup. Table 1). Organoids containing *Nf1* mutations, such as TN, commonly resulted in poorly differentiated mesenchymal-like (sarcomatoid) neoplasms (Fig. 4C). Such neoplasms were largely composed of tightly arranged spindle and polygonal cells marked by a medium to high proliferation index according to nuclear Ki67 immunostaining, and detected mitotic figures. Such cells contained low to moderate of amount of cytoplasmic KRT8 and nuclear PAX8, indicating an epithelial tumor that developed mesenchymal features. Organoids with *Map2k4* mutations predominantly formed papillary-glandular carcinomas with slit-like spaces forming labyrinthine patterns typical for HGSC (8/17 vs 3/38 for other mutations Fisher’s P=0.0019). Cells of such neoplasms consistently expressed KRT8 and PAX8 and had regionally variable proliferative indices and high-grade nuclear atypia. Remaining neoplasms exhibited both mesenchymal and papillary-glandular patterns of growth. All neoplasms invaded underlying striated muscles and upper levels of dermis at the place of organoid transplantation. Additionally, we noticed that *Nf1* mutation-containing tumors exhibiting spindle cells were derived from organoids with well-defined lumens, while tumors with glandular patterns arose from organoids with more blob-like structures (Fig. 3H). These findings relate specific genetic alterations to distinct features of HGSC.

### Tumor transcriptomes align with pathology

Transcriptomic analyses revealed that mesenchymal and papillary tumors grouped well in PCA (Fig. 5A). When compared to human tumor samples, mesenchymal tumors were consistent with human mesenchymal phenotypes whereas papillary tumors eluded decisive subtype classification (50) (Fig. 5B). To further elucidate differences between these two phenotypes, we compared *Trp53/Map2k4/Pten* to *Trp53/Nf1/Pten* tumors. GSEA indicated that the former were epithelial-like while the latter was characterized by mitosis and proliferation hallmarks (Sup. Fig. 8AB). We also noticed that TMP tumors have downregulated proteolysis activity, which suggests a genotypic preference of metastasis avoidance in *Map2k4* mutated tumors (Fig. 5C).

**Figure 5.**
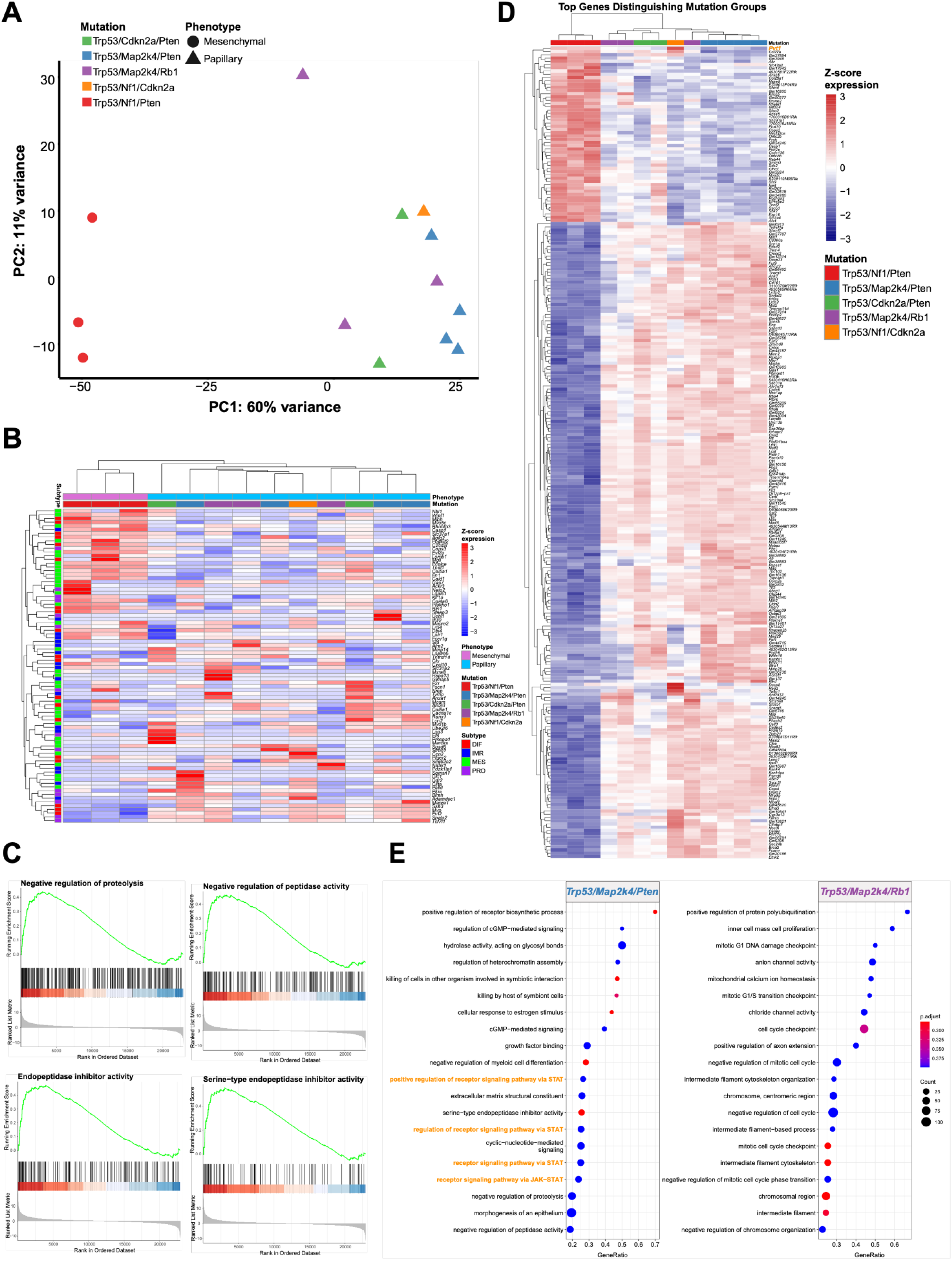
Transcriptomic analysis of tumors. A) PCA analysis of mesenchymal and papillary tumors. B) Alignments of mesenchymal and papillary tumors against human gene expression subtype signatures (50). C) GSEA analysis of Trp53/Map2k4/Pten vs Trp53/Nf1/Pten tumors. D) Top 250 transcriptomic differences from a likelihood ratio test (LRT) on genotype. Pvt1 gene is highlighted at top in orange. E) Comparison of GSEA hallmarks in Trp53/Map2k4/Pten and Trp53/Map2k4/Rb1 tumors.

We next explored whether specific gene disruptions guided differential tumor presentations. In all *Nf1* mutant tumors presenting with either papillary or mesenchymal phenotypes, there was upregulation of *Pvt1* (a lncRNA regulated by *Myc*) (Figure 5D). This is consistent with published human data where *Pvt1* is one of the top amplified genes (cBioPortal) and increased expression is associated with poor survival outcome (Sup. Fig. 8C). Within the papillary subgroup, those with TMP mutations had upregulated JAK-STAT signaling compared to those with TMR mutations. Together, these observations highlight that mutational signatures must be considered in addition to pathology phenotypes for patient treatment (Fig. 5E).

### Mutation combinations influence responses to common HGSC therapeutics

Due to differences in presentation of *Nf1* and *Map2k4* mutations at the organoid and tumor level, we tested how organoids bearing these mutations would respond to common HGSC treatments. We performed dose-response survivability assays with carboplatin (a cisplatin analog causing DNA crosslinks), gemcitabine (a base analog that inhibits DNA replication), niraparib (PARP inhibitor), paclitaxel (a microtubule inhibitor that blocks mitosis), and trametinib (MEK inhibitor), with or without ROCK (Rho-associated protein kinase) inhibitor (ROCKi; a disruptor of cytoskeletal function).

Removing ROCKi from the culture media increased resistance (increased in the inhibitory concentration) of transformed organoids to carboplatin (180%), gemcitabine (122%), and niraparib (133%), while paclitaxel sensitivity remained unchanged and trametinib became more effective (64%) (Sup. Fig. 7A). *Nf1* or *Map2k4* deficient organoids treated with carboplatin or gemcitabine with or without ROCKi did not have any significant differences (Fig. 6AB). *Map2k4* mutants were generally more sensitive to paclitaxel and human data suggests a correlation between low *MAP2K4* expression and improved survival when treated with taxols (paclitaxel). In line with our transcriptomic findings, *PVT1* overexpression (as observed in *MYC*-amplified tumors and in our *Nf1*-mutated tumors) is associated with paclitaxel resistance and poorer survival outcomes (Fig. 6D, Sup. Fig. 7BC). Trametinib, a treatment for neurofibromatosis, caused *Map2k4*^mut^ to be more resistant than *Nf1*^mut^, however these effects were abolished upon ROCKi co-treatments (Fig. 6E). Interestingly, TNM cells were unaffected by trametinib, contrary to expectations, as we expected *Map2k4* mutations to act similarly to *Nf1* mutations (Fig. 6E), and that inhibition of both the RAS and JNK arms of the RTK signaling pathways are effective in inhibiting cancer cell growth (51–53). These differences could be due to differences in p53 status or the culture conditions that prevent apoptosis. Removal of ROCKi increased resistance in *Map2k4* mutants, while no correlation between *Map2k4* mutations and trametinib sensitivity was seen with ROCKi (Fig. 6E), suggesting that ROCKi may play a role in promoting cell survival. Differences in *Nf1* and *Map2k4* mutation responses to chemotherapeutics, particularly the high resistance in TNM combinations, demonstrate the need to study specific mutational perturbations.

**Figure 6.**
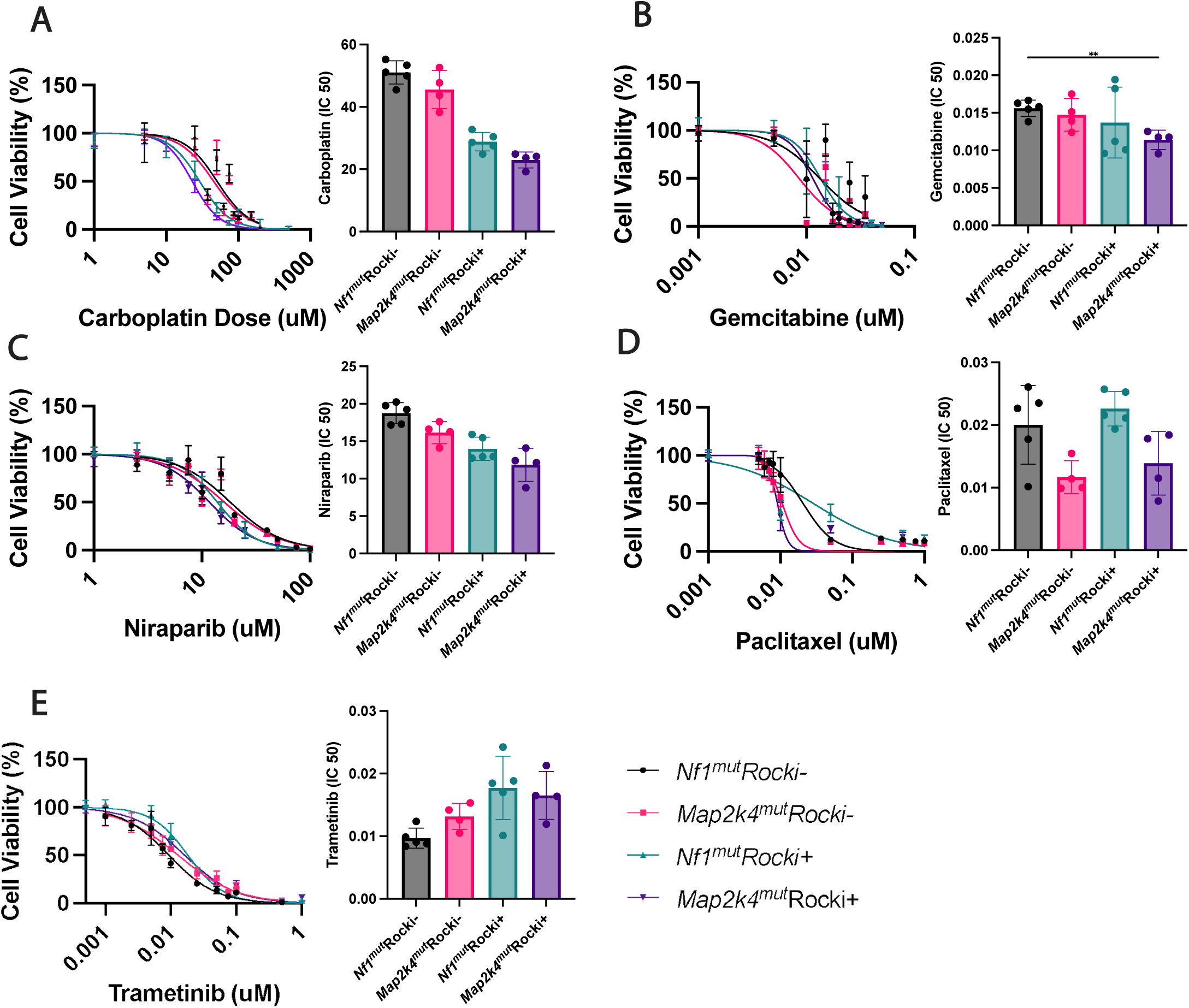
Impacts of drug treatments to growth of organoids bearing *Map2k4* or *Nf1* mutations. A-E) Dose response curves to indicated drugs in the presence or absence of ROCKi are plotted. One-way anova, Brown-Forsythe and Welch Tests were performed to assess differences in sensitivity and resistance. Two independent experiments were conducted per mutation combination in triplicate. One-way anova was used to assess significance. P-value < * = 0.05, <** = 0.01, <*** = 0.001, <****= 0.0001.

## DISCUSSION

HGSC presents as a heterogeneous solid tumor that is genetically unstable with multiple clonal lineages (54,55). These aspects have made treating HGSC especially challenging in stage III-IV HGSC, where 70% of patients relapse within 3 years (56). Early evidence suggests that HGSC does not follow a typical tumor growth trajectory; instead, dissemination occurs as a single rapid wave late in the tumor’s evolutionary trajectory (57). Therefore, tumor initiation, progress, and presentation will be pivotal for developing diagnostic tools at earlier stages and improving patient outcomes.

Whereas HGSC can originate from both OSE and TE cells it is unknown if they utilize the same driver combinations. In our TE organoid screen, we used the same CRISPR KO mini library and infection conditions as we used in a prior screen of OSE (45). The transformation rates were similar: 0.28% for TE and 0.21% for OSE (45). Both OSE and TE transformation involved key HGSC drivers (*Trp53*, *Cdkn2a*, *Rb1*, *Pten*) (Sup. Fig. 8A) (8,9,11,12,15,45), but TE transformation was further enhanced by *Nf1*, *Map2k4*, *Brca2*, and *Brca1* mutation, while OSE transformation was enhanced by *Fancd2*, *Wwox*, *Gabra6*, and *Fat3* mutations (Sup. Fig. 8A). Notably, combinations like *TNP+Fancd2* or *TMC+Wwox* did not increase TE organoid transformation beyond their core triple mutant counterparts (Sup. Fig. 4C, D). Pathway analysis suggests that genes like *Wwox, Gabra6*, and *Fat3* may regulate the MAPK and AKT pathways, while *Fancd2* is involved in DNA repair and interacts with BRCA1/2 (Fig. 7) (58–68). Despite some gene-specific differences, there may be shared altered pathways during malignant transformation. Other studies have compared the transformation potential of OSE and TE cells in organoid models and found differential results with respect to tumorigenic potential (10,11). However, OSE cells are flat cuboidal cells that exist as a single monolayer surrounding the ovary whereas TE cells exist in a pseudo-stratified structure that requires a 3D space. To this extent, our screening methods to assess transformation potential by adherence-independent growth (OSE) and aberrant morphology (TE) are more suitable approaches for maintaining cells in their native state. In summary, *Fancd2, Wwox, Gabra6,* and *Fat3* appear OSE-specific, while *Nf1, Map2k4, Brca2*, and *Brca1* are TE-specific, though these findings need further validation in direct comparisons.

**Figure 7.**
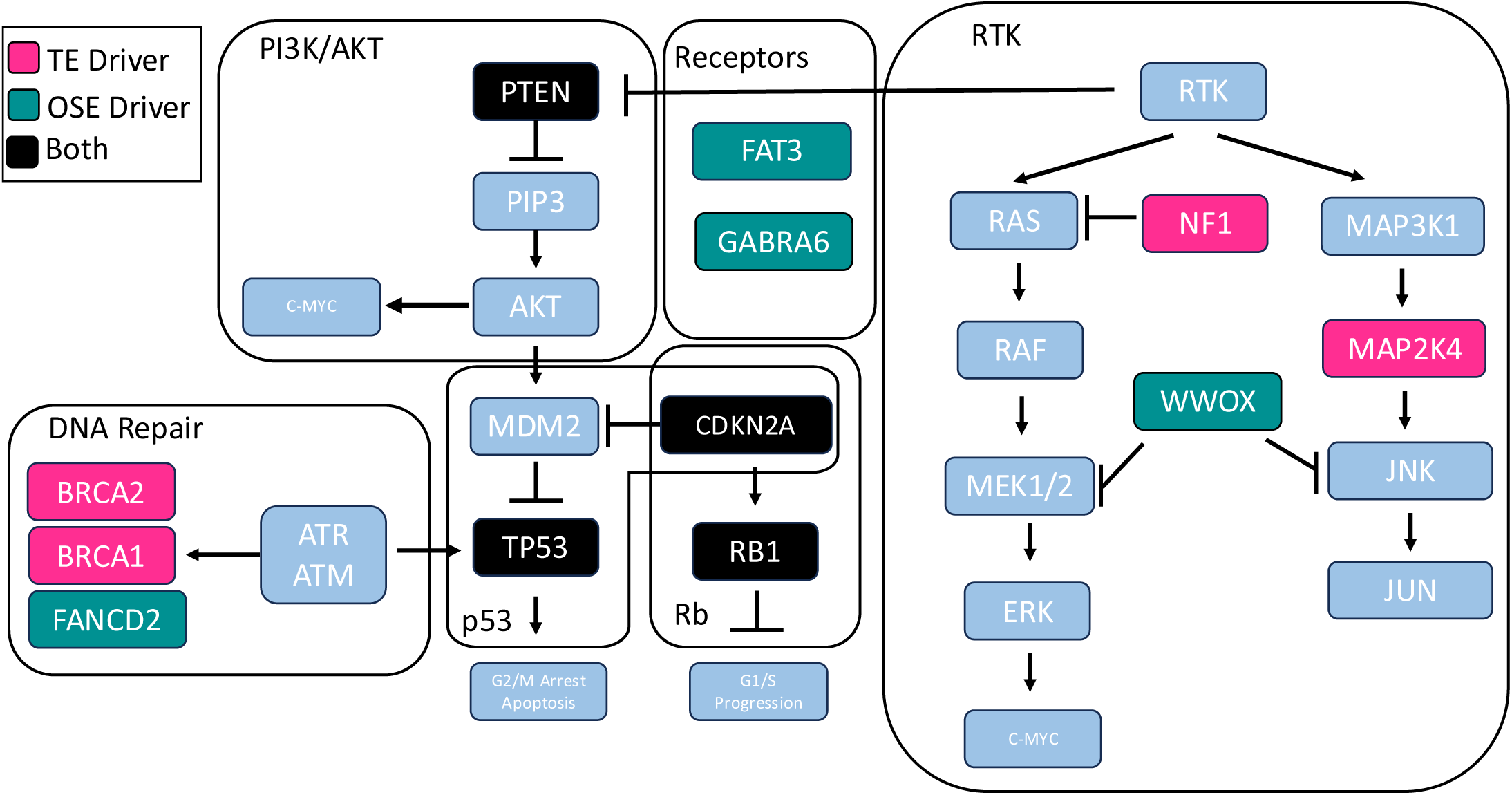
Pathways of gene drivers identified in OSE and TE screens.

Of the seven transformation driver genes identified in our screen, *Map2k4* was the most intriguing. Encoding MKK4 in the JNK pathway, *Map2k4* phosphorylates JNK or p38 in a context-dependent manner (46,52,53,69). Evolutionary analyses of human HGSC suggest that aberrant MAPK signaling is associated with an early evolutionary state, marked by fewer subclones, diploid cancers, and better therapy response (70). Further tumor trajectory analysis implicates *MAP2K4* mutations in metastasis, particularly in the dissemination from the ovary to the omentum and ascites (57). While *Map2k4* overexpression can reduce tumor metastasis (71,72), our data show enhanced organoid transformation when co-mutated with *Nf1, Cdkn2a, Rb1, Pten*, or *Brca1*, though not all combinations formed tumors within 200 days (Sup. Table 1). This suggests that aberrant organoid formation may reflect metastatic potential or tumor subtype presentation, rather than just carcinogenesis (72). Our findings experimentally establish *Map2k4* as a tumor suppressor and show that tumors containing *Map2k4* genetic alterations have more differentiated papillary phenotypes typical for HGSC, suggesting this gene as a potentially useful target for intervention. It also opens the possibility for deriving more accurate mouse models of HGSC involving inactivation of *Map2k4*.

Modern precision cancer treatment has been revolutionized by considering the key genes involved in a person’s cancer. When cancer cells survive and gain mutations that promote resistance following first-line treatments, patient survival is often poor. For example, platinum-based therapeutics are initially highly effective but become less successful with each round of treatment (73). Therefore, integrating genetics with treatment modalities is integral for improving patient outcomes. The organoid platform enables facile screening of anticancer agents that are most effective against transformed cells bearing various mutant combinations.

Recent research on HGSC treatment has focused on how genetic perturbations influence responses to common therapies (carboplatin, gemcitabine, niraparib, paclitaxel, trametinib). Other studies have shown that differences in cell of origin and chemoresponse exist (10,11). For instance, WT OSE and TE organoids exhibited differential responses to HGSC treatments, highlighting the impact of culture conditions on drug efficacy. One group has shown that genes regulating similar pathways, such as RAS, have differences in response to common therapeutics that are gene context-dependent, such as *MYC^OE^, KRAS^OE^,* or *NF1^-/-^* (44).

We found that *Map2k4* mutants were particularly sensitive to paclitaxel treatment, providing a potentially new avenue of treatment when considering JNK pathway-altered cancers, primarily those implicating *Map2k4* and its downstream targets. ROCKi did not impact the outcomes of paclitaxel treatment. We also observed that *Rb1/Cdkn2a* mutant cells responded similarly to gemcitabine regardless of ROCKi addition, though there were two subgroups: one more sensitive in the presence of ROCKi, the other less so. Generally, *Pten* and *Map2k4* mutations increased sensitivity, while *Nf1* mutations contributed to resistance in the presence of ROCKi. Previous studies have shown that overexpression of *Ccne1* enhances gemcitabine sensitivity (44), and we found *Trp53/Rb1* mutants to be among the most resistant, with *Nf1* mutations maintaining resistance and *Map2k4* mutations increasing sensitivity. Other groups have shown that *Pten* mutations tend to drive resistances when compared to *Nf1* (44), but we found the opposite in our system. Altogether this may be due to differences in culture conditions or assay design. Nonetheless this implores the need to consider the whole tumor and identifying dominant mutations that are important for diagnosis and treatment of patients.

ROCKi (e.g., Fasudil) is emerging as a promising tool in combination therapy due to its roles in cytoskeletal regulation, cell migration, proliferation, survival, and metastasis (74,75). Previous studies have shown its potential to enhance chemo-sensitivity in resistant cancers, such as pancreatic cancer when combined with gemcitabine (76). Our data support this, showing that ROCKi treatment with gemcitabine significantly reduced transformed organoid survival compared to untreated controls (Sup. Fig. 6A). ROCKi also increased sensitivity to carboplatin and niraparib but conferred resistance to trametinib (Sup. Fig. 7A). Notably, transformed TE cells with *Nf1* mutations showed more resistance to paclitaxel than those with *Map2k4* mutations, with no significant difference between ROCKi –treated and untreated conditions. However, *Map2k4* mutations were more resistant to trametinib without ROCKi. While other groups have shown that RAS-driven cancers containing *Map2k4* mutations are more sensitive to trametinib treatment, our data shows the opposite and our most resistant combination was TNM (51–53). These findings suggest that ROCKi could enhance treatment efficacy for specific mutations, offering new therapeutic possibilities.

While previous groups have shown that *TNP* mutations readily form tumors (44), our study goes further by detailing the specific impact of these mutations on tumor progression (Fig. 4). We also show that organoid metrics of transformation and diameter correlate with patient survival, a previously unreported correlation. Additionally, we report that *Nf1* mutants preferentially form mesenchymal tumors while *Map2k4* mutants preferentially form papillary tumors (Fig. 4). *Nf1* mutations also upregulate *Pvt1*, a long non-coding RNA (lncRNA) that MYC can bind and regulate (77). Other lncRNA’s such as MALAT-1 and DANCR have also been implicated in ovarian cancer via c-MYC activity (78). Additionally, combination therapy of c-MYC inhibitors and paclitaxel have been shown to increase sensitivity to previously paclitaxel resistant cell lines (79). This observation suggest the potential utility of developing new mouse models focused on deficiencies of *Map2k4, Nf1* and certain lncRNAs as a potential avenue for exploring HGSC treatments.

Our previous OSE screen showed that stem cells are more prone to transformation than non-stem cells (45). This study was consistent with previous findings that stem/progenitor OSE cells are preferentially susceptible to transformation by inactivation of tumor suppressor genes *Trp53* and *Rb1* (8). Our recent studies have shown that, unlike OSE, inactivation the same genes in the TE lead to preferential HGSC formation from pre-ciliated cells (15). Thus, specific cell state of differentiation may play a very significant role in cell susceptibility to transformation.

Future studies are required to test if TE cells in different differentiation states may also have different propensity for transformation and chemoresistance as a function of specific HGSC drivers. We recently described TE cell lineage hierarchy and showed distinct expression patterns of *Map2k4* and *Nf1* along TE cell lineage trajectories (15). Specifically, *Map2k4* expressed in secretory and ciliated cell lineages, while *Nf1* expression was mainly limited to stem/progenitor cells and pre-ciliogenic cells. Our current study demonstrated that *Map2k4* mutations tend to form papillary-glandular carcinomas, while *Nf1* mutations resulted in mesenchymal phenotype. It will be of particular interest to clarify if expression of *Map2k4* in the secretory TE cell lineage pre-determines preferential papillary-glandular phenotype and/or sensitivity to paclitaxel and resistance to trametinib by resulting tumors.

Immunocytological and functional assays reported here and elsewhere (10, 11, 44) revealed that TE organoids contain stem/progenitor cells (PAX8+), ciliated cells (FOXJ1+, acetylated alpha-tubulin+) and secretory cells (OVGP1+). Thus, future organoid studies in conjunction with combinatorial mutagenesis will facilitate identification of the most vulnerable cancer-prone cell states, their genetic drivers and chemoresistance traits. Additionally, our platform is amenable to modeling of amplification/overexpression events using CRISPRa (80) and assessing how gene combinations respond to therapies. The high throughput and speed of our platform has allowed us to identify novel genetic drivers and optimal treatment strategies and can be further translated into human models to quickly assess patients for the most beneficial treatment outcomes.

## METHODS

### Experimental animals

NOD.Cg-*Prkdc*^scid^ *Il2rg*^tm1wjl/^SzJ (NSG) mice (Stock number 005557 were obtained from The Jackson Laboratory (Bar Harbor, ME, USA). The number of animals used in every experiment is indicated as biological replicates in Fig. legends and supplementary tables. Animals were euthanized if they became moribund or developed tumors over 1 cm in diameter. All the experiments and maintenance of the mice were conducted under protocols approved by the Cornell University Institutional Laboratory Animal Use and Care Committee (protocols numbers 2000-0116, 2001-0072 and 2004-0038). Mice were housed within a 10/14 light cycle. The lights came on at 5 a.m. and went off at 7 p.m., the humidity ranged from 30 – 70 %, and the ambient temperature was kept at 72°F +/-2°F.

### TE organoid derivation and culture

TE Organoids were cultured and derived as previously described (15). TE cells were isolated and cultured into organoids from 2-3 month old *Gt(ROSA)26Sor^tm9(CAG-tdTomato)Hze^* (*Rosa-loxp-stop-loxp-*tdTomato/Ai9, Stock number 007909 Jackson Laboratory) were sacrificed, oviducts isolated and dispersed into single cells with collagenase/dispase. Cells were rescued with 20% FBS containing media and plated as rims with matrigel (Corning #356231) at a density of 10kcells/100uL of matrigel per well in a 24w plate. Organoids were then expanded and cultured for at least 5 passages before experiments were performed. Organoids were passaged on day 7 unless otherwise specified. To passage organoids, media was aspirated from the wells, 300uL of cultrex organoid harvesting solution (R&D 3700-100-01) was added to each well and vigorously pipeted to disrupt matrigel. Dissolved matrigel was then transferred to a 15mL conical and allowed to rest on ice for 1hour. Organoids were then spun down at 600xg for 5 minutes at 4C. If matrigel is still present, more cultrex organoid harvesting media was added and incubated for 30 minutes followed by centrifugation as described above. Once all matrigel is gone, supernatant was removed and 1mL of Trypsin EDTA 0.25% (Gibco 25200072) was added and incubated at 37C. Every 5 minutes organoids were disrupted into single cells with a P1000 pipet (vigorous pipetting at least 50 times up and down). This step was repeated 3 times for a total of 15 minutes. 20% FBS containing media was then added and cells were pelleted as described above. Cells were then resuspended in TE media and counted to be passaged as described above.

### Lentiviral production and transduction

Lentivirus were produced in 293T cells and titer determined via limiting dilutions of virus in TE organoid cells as previously described (45). Briefly, 293T cells were seeded into 10cm dishes before LentiCRISPRv2 (Addgene #52961), PMD.2G (Addgene #12259) and PSPAX2 (Addgene #12260) were transfected using Transit-LT1 transfection reagent (Mirus bio, MIR2306) via manufacturer instructions. Virus was collected at 48 and 72 hours post transfection. Lenti-X Concentrator (Takara, 631231) was used as instructed to concentrate virus. Virus was then resuspended in TE organoid media with polybrene (Sigma, TR-1003-G) and frozen at –80C until use. LentiCRISPRv2-GFP (Addgene #82416) reporter plasmid was used to evaluate the quality of virus produced. To titer the virus, 5K TE organoid cells were subjected to limiting dilutions of virus in a 96 well plate and spinoculated at 600xg for 1hour at 37C as previously described (44). Lentiviral transduction continued for 6 hours at 37C in a cell incubator before cells were collected, mixed with matrigel and plated. After 48 hours 2ug/mL of puromycin was used to select for properly infected cells. Infection percent was evaluated by determining the number of surviving cells in treated versus non treated samples. The multiplicity of infection was calculated by using a survival percent below 30% to exclude double infection events. These values were then multiplied and averaged to determine an MOI of 7 (45).

### Screen and validation

Day 7 TE organoids were broken into single cell suspensions as described above and infected with the minilibrary at an MOI of 7 and plated at a density of 5K cells per 25uL Matrigel droplet. 48 hours post infection, cells were treated with puromycin (2ug/mL, Sigma, A1113803) and at 96 hours media was changed to contain 10uM Nutlin-3A (SelleckChem, S8059). Media was changed every 2-3 days containing 10uM Nutlin-3A until day 14 where organoid transformation was assessed. Mutant organoids were then released from Matrigel with a P1000 wide bore tip in cultrex organoid harvesting media and allowed to settle by gravity. Supernatent was aspirated and replaced with more cultrex organoid harvesting media and allowed to settle by gravity. Organoids were then released onto a poly-hema (Sigma, P3932) coated tissue culture plate, picked as whole organoids, expanded in 2D culture, genomic DNA harvested with GeneJET Genomic DNA Purification Kit (Thermo Fisher, K0722), and PCR amplification with the following primers: CTTGGCTTTATATATCTTGTGGAAAGG and CGACTCGGTGCCACTTT was performed with Illumina overhangs also added to each primer. Indexing reactions were performed to barcode each individual colony and pooled. 300bp paired end sequencing was performed using Illumina Miseq to detect gRNA presence in a particular clone at a read depth of 4 million. Each sequenced clone was aligned with a custom built library containing each of the gRNA’s in the screen to identify “hits,” where genome integration had taken place as described previously (45,80).

After identification of key combinations that result in TE organoid transformation, targeted lentiviral libraries were prepared and single cell suspensions from TE organoids were infected with the libraries at a minimum MOI of 7. Cells were then plated, treated with both puromycin and Nutlin-3A as described above. Organoid transformation rate was determined by counting the total number of aberrant organoids and dividing that by the total number of organoids formed after 14 days of culture. Images were acquired on a Zeiss Axiovert microscope.

### Organoid preparation for histological evaluation

Organoids were released with organoid harvesting solution as described above. Organoids were placed on ice for 1 hour to release and allowed to settle to the tube bottom before aspirating the media without disturbing the pellet. Organoids were then washed with 10mL of ice cold 1X PBS to remove any residual Matrigel. Organoids were allowed to settle by gravity, the PBS was aspirated, then fixed in 4% PFA (diluted in PBS) on ice for 1 hour. Organoids were then washed twice with ice cold 1X PBS, then mixed with an equal volume of HistoGel (Epredia, HG4000012) preheated to 65°C and dispensed into dome-shaped molds for embedding. HistoGel molds set for 10 minutes prior to standard tissue processing, paraffin embedding, and preparation of 4-μm-thick tissue sections followed by immunohistochemistry (15,81).

### Mouse tumorigenic studies

According to previous results (10), frequency of tumor formation by mutant TE organoids was not impacted by site (s.c. vs, orthotopic) of organoid transplantation. Targeted library preparation and TE organoid infections were performed as described above. After passaging to expand cells, day 14 organoids were then disrupted into single cells and 5×10^5^ cells were mixed with 100uL of 1X PBS (Gibco 10010023) and Matrigel (Corning #356231) at a 1:1 ratio and injected subcutaneously in the flanks of NSG mice. All mice received bilateral injections and were monitored weekly for tumor formation, and growth of tumors. Mice were sacrificed when tumors reached 1 cm in diameter.

### Drug Assays

Organoids were grown and dispersed into single cells as described above. Cells were diluted to a concentration of 100 cells per uL of Matrigel. 10uL of Matrigel containing cells were then seeded per well into the bottom of a black walled clear bottom 96 well plate (Corning #3603) and 100uL of media was added. 1 day after plating, cells were treated with the concentrations of drugs, media was changed on day 3 and on day 5 media was removed and 100uL of 3D Cell titer glo (Promega # G9683) was added for 1hr at room temperature and read on a biotek plate reader (10). Prism 10 was used to fit graphs and interpret IC50 values with the “[inhibitor] vs. response-variable slope (normalized)” option.

### Immunohistochemistry and image analysis

For paraffin embedding, organoids and tissues were fixed in 4% paraformaldehyde overnight at 4°C followed by standard tissue processing, paraffin embedding and preparation of 4 μm-thick tissue sections. For immunohistochemistry, antigen retrieval was performed by incubation of deparaffinized and rehydrated tissue sections in boiling 10 mM sodium citrate buffer (pH 6.0) for 10 minutes. The primary antibodies were incubated either at 4°C for overnight or at room temperature for 2 hours, followed by incubation with secondary biotinylated antibodies (dilution 1:200, 45 minutes, at room temperature, RT). Modified Elite avidin-biotin peroxidase (ABC) technique (Vector Laboratories, Burlingame, CA, USA; pk-6100) was performed at room temperature for 30 minutes. Hematoxylin was used as the counterstain. All primary and secondary antibodies used for immunostaining are listed in Supplementary Table 2. For quantitative studies, sections were scanned by ScanScope CS2 or FL (Leica Biosystems, Vista, CA) with a 40× objective, followed by the analysis with the ImageJ software (National Institutes of Health, Bethesda, MD, USA).

### RNA isolation, sequencing and analysis

Tumor samples were collected and immediately flash frozen in liquid nitrogen. All RNA samples were prepared from tumors using the Zymo *Quick* RNA extraction kit (R1054) and immediately sent for quality control, mRNA library preparation (poly A enrichment) and NovaSeq X Plus Series (PE150) (6.00G raw data per sample). After quality analysis with FastQC and processing read lengths with trimmomatic (v0.39), reads were aligned to mm10 reference genome with hisat (v2.2.1) and SAMtools (v1.21). Count matrices for each sample were generated using the subread (v.2.0.8) package. The count matrices were analyzed and visualized using R (v4.05) with DEseq2 (v1.30.1), biomaRt (v2.46.3), pheatmap (v1.0.12), and enrichplot (v1.10.2). The RNA-seq datasets can be found at accession number xxxxxxx.

## Statistical analyses

Statistical comparisons were performed using a two-tailed Fisher’s exact test with InStat 3 and Prism 6 software (GraphPad Software Inc., La Jolla, CA, USA). Prism 10 software was used to perform one-way anova, IC50 calculations, welch’s two tailed t-test. cBioportal was used to perform analysis of human data. χ^2^ analysis was performed in excel. KM Plotter was used on ovarian cancer datasets.

## Acknowledgements

This work was supported by NIH grants (CA182413, CA260115 and CA248524) to AYN, Ovarian Cancer Research Fund grant (327516) to AYN, Sandra Atlas Bass Endowment for Cancer Research to AYN and JCS, and the NSF Graduate Research Fellowship Program (GRFP) awarded to CQR (DGE-2139899).

## Author contributions

DJP, JCS and AYN designed experiments; DJP, CQR, TME, CSA and APA performed experiments and performed pathological evaluations; AFN, JCS, AYN and RJY provided resources or funding; DJP wrote the paper; DJP, JCS, AYN revised and edited the paper.

