## Supplemental for "Combinatorial organoid mutagenesis screen reveals gene constellations driving malignant transformation, pathology and chemosensitivity in high-grade serous ovarian carcinoma"

Phuong et al

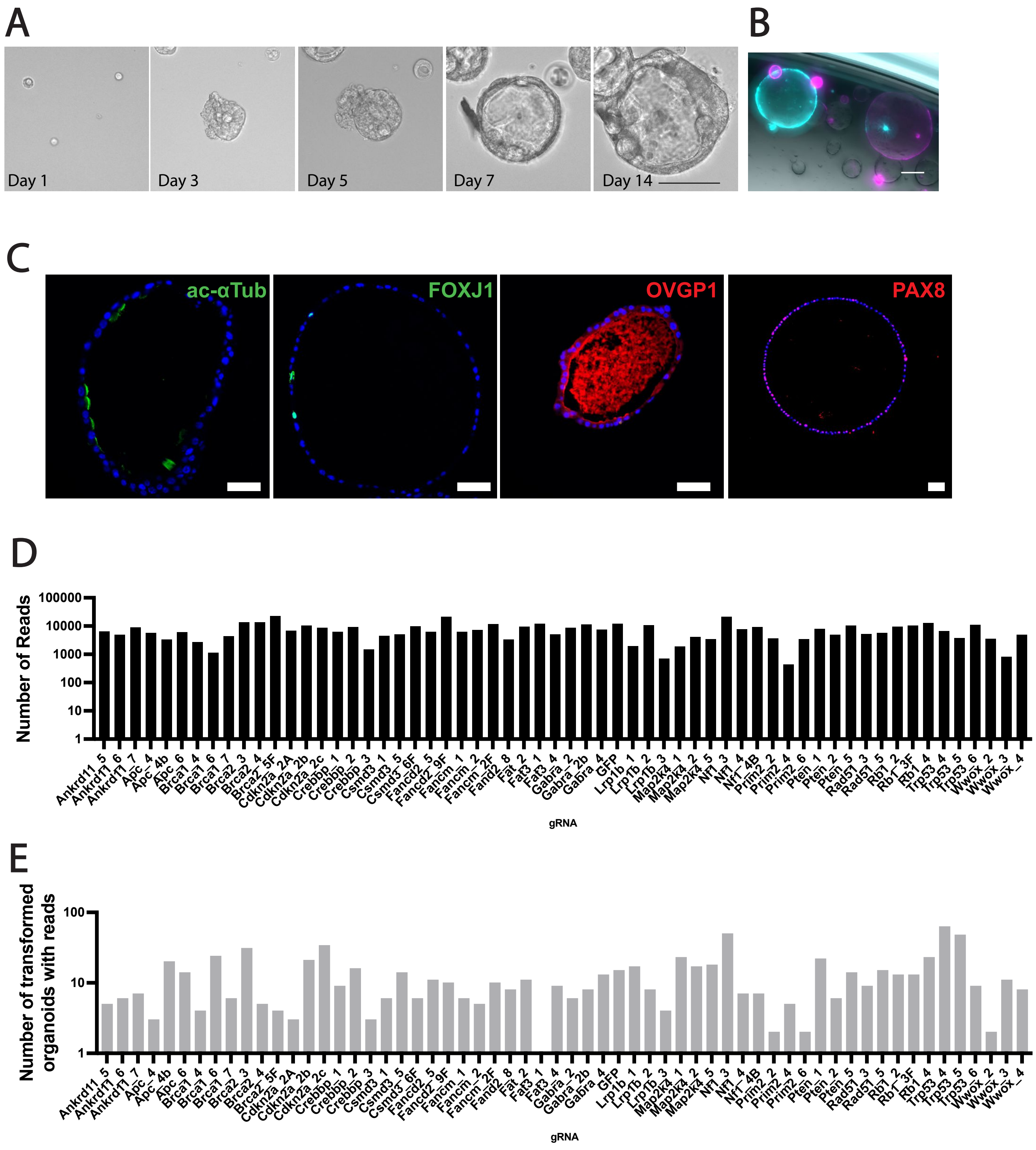


**Supplementary Figure 1. Validation of TE organoids and minilibrary for CRISPR screening**. A) TE Organoids grow and self organize. Scale bar = 100µm. B) TE Organoids express either mCherry (magenta) or GFP (teal) after lentiviral infection with a lack of signal co-localization. Scale bar = 100µm. C) TE Organoids express canonical TE markers. Scale bar = 50µm. D) LentiCRISPR mini-library distribution post infection in 293T cells. E) gRNA presence in picked and expanded clonal aberrant organoids from screen.

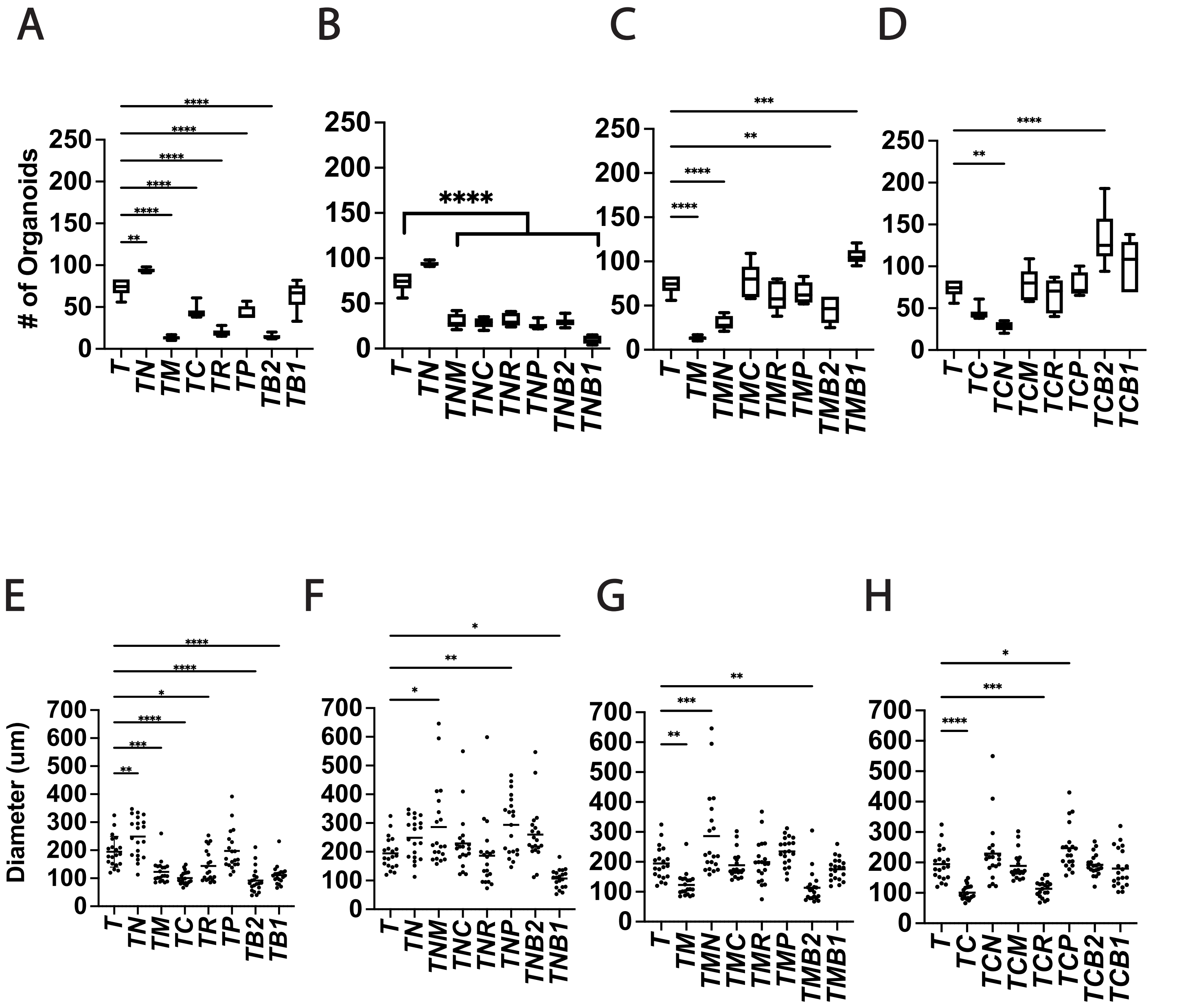


**Supplementary Figure 2**. **Validation of organoid formation and diameter from combinations identified in screen.** ABCD) Formation of organoids with different combinations. N = 6. EFGH) Organoid diameters. n= 20 organoids. One-way anova was performed to assess significance. P-value < * = 0.05, <** = 0.01, <*** = 0.001, <****= 0.0001. *Trp53*, abbreviated ("T"), *Nf1* (N), *Map2k4* (M), *Cdkn2a* (C), *Rb1* (R), *Pten* (P), *Brca2* (B2) and *Brca1* (B1).

A

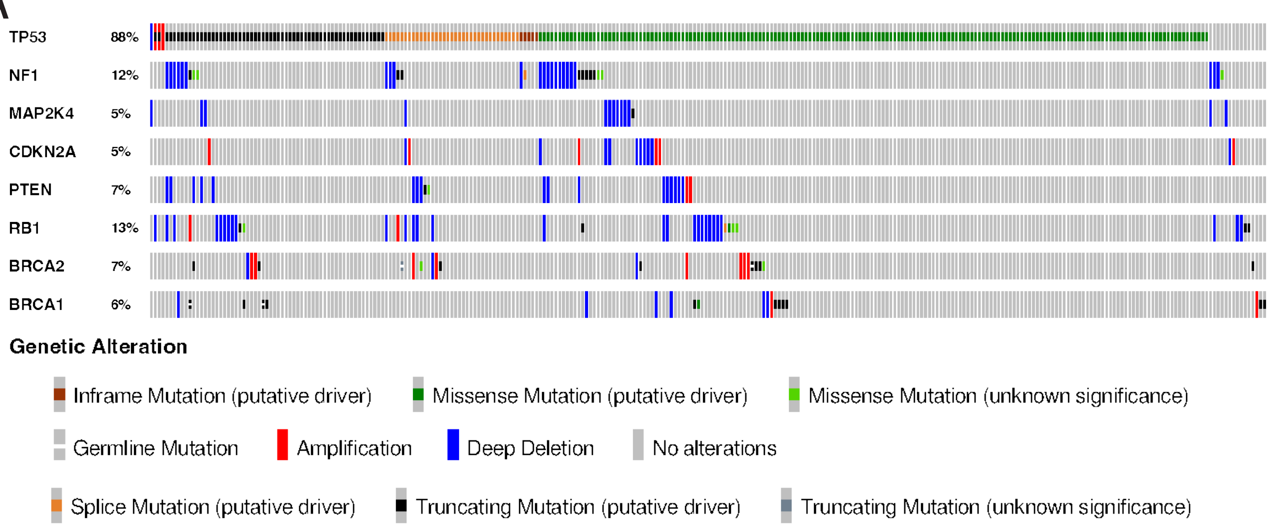


B

| Gene A | Gene B | Samples with neither | Samples with Gene | Samples with Gene | Samples with both | p-Value | q-Value | Tendency |
|---|---|---|---|---|---|---|---|---|
| *MAP2K4* | *CDKN2A* | 251 | 8 | 11 | 3 | 0.014 | 0.226 | CO |
| *NF1* | *PTEN* | 225 | 27 | 15 | 6 | 0.028 | 0.226 | CO |
| *RB1* | *PTEN* | 224 | 28 | 15 | 6 | 0.032 | 0.226 | CO |
| *PTEN* | *BRCA2* | 235 | 17 | 17 | 4 | 0.065 | 0.342 | CO |
| *NF1* | *RB1* | 213 | 26 | 27 | 7 | 0.153 | 0.644 | CO |
| *RB1* | *BRCA1* | 226 | 30 | 13 | 4 | 0.243 | 0.851 | CO |
| *CDKN2A* | *BRCA2* | 240 | 12 | 19 | 2 | 0.293 | 0.88 | CO |
| *NF1* | *MAP2K4* | 229 | 33 | 11 | 0 | 0.371 | 0.973 | ME |
| *NF1* | *BRCA1* | 226 | 30 | 14 | 3 | 0.442 | 1 | CO |
| *MAP2K4* | *PTEN* | 242 | 10 | 20 | 1 | 0.593 | 1 | CO |
| *CDKN2A* | *BRCA1* | 243 | 13 | 16 | 1 | 0.603 | 1 | CO |
| *NF1* | *CDKN2A* | 228 | 31 | 12 | 2 | 0.68 | 1 | CO |
| *RB1* | *BRCA2* | 220 | 32 | 19 | 2 | 1 | 1 | ME |
| *MAP2K4* | *BRCA2* | 241 | 11 | 21 | 0 | 1 | 1 | ME |
| *PTEN* | *BRCA1* | 236 | 20 | 16 | 1 | 1 | 1 | ME |
| *BRCA2* | *BRCA1* | 236 | 20 | 16 | 1 | 1 | 1 | ME |
| *CDKN2A* | *PTEN* | 239 | 13 | 20 | 1 | 1 | 1 | ME |
| *CDKN2A* | *RB1* | 226 | 13 | 33 | 1 | 1 | 1 | ME |
| *MAP2K4* | *RB1* | 229 | 10 | 33 | 1 | 1 | 1 | ME |
| *NF1* | *BRCA2* | 221 | 31 | 19 | 2 | 1 | 1 | ME |
| *MAP2K4* | *BRCA1* | 245 | 11 | 17 | 0 | 1 | 1 | ME |

**Supplementary Figure 3**. **Analysis of genetic alterations in human HGSC.** A) OncoPrint (from cBio Portal; Firehose legacy) of human HGSC genetic alterations of selected tumor suppressor genes. B) Co-occurrence of gene drivers identified in screen as seen in human HGSC (only samples containing a *TP53* mutation were assessed). CO = Co-occurrence, ME = Mutual Exclusivity.

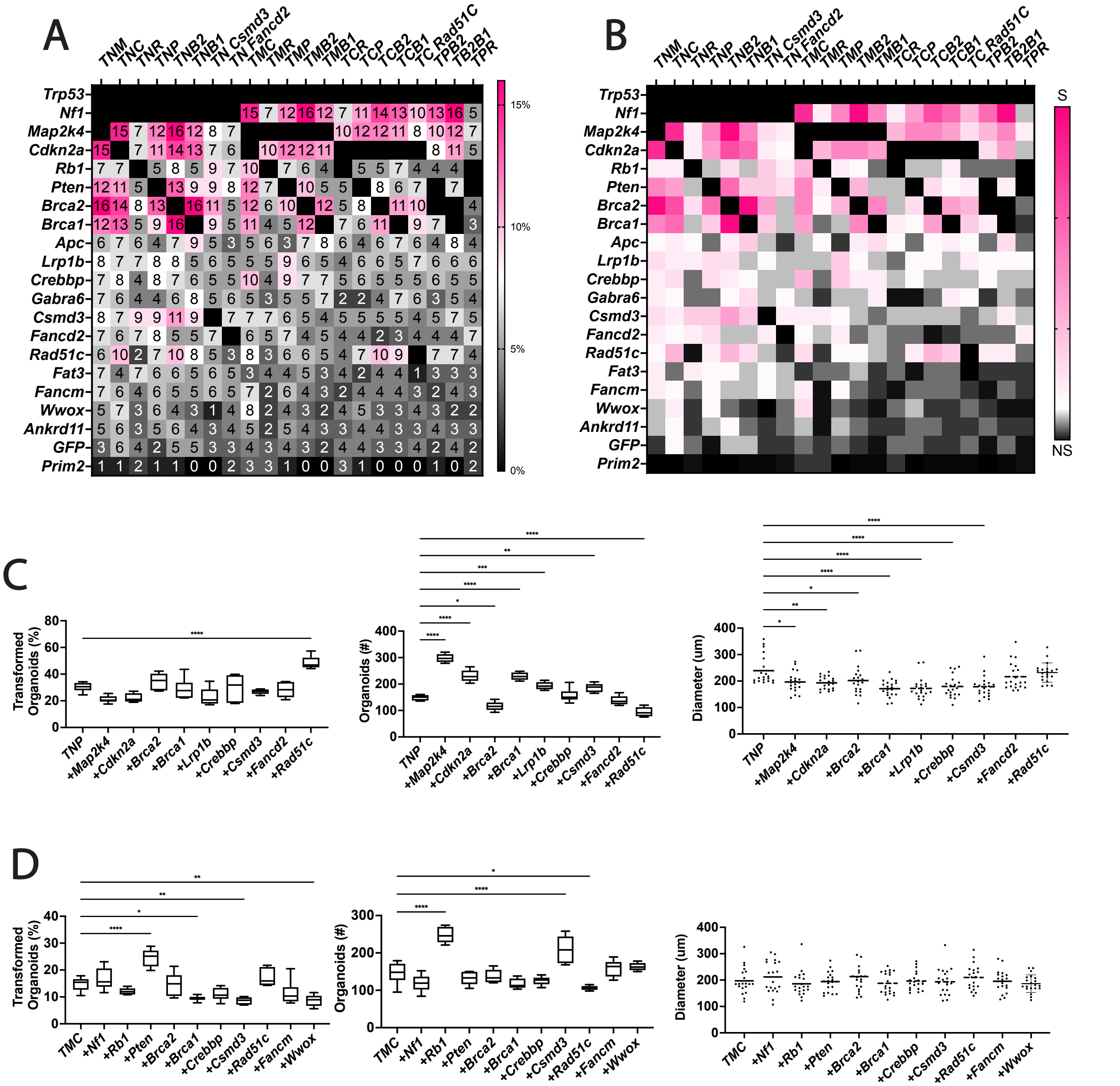


**Supplementary Figure 4**. **Analysis and validation of quadruple mutant combinations.** A) Starting with the most overrepresented triple combinations, the most significant quadruple mutations are plotted. B) $\chi^2$ analysis of significant TE triple quadruple mutants, DF, 19, P<0.05%. S = Significant; NS = Not Significant. C) Transformed organoids in a *Trp53*$^{-/-}$*Nf1*$^{mut}$*Pten*$^{mut}$ deficient background. D) Organoid formation efficiency. E) Organoid size. F) Transformed organoids in a *Trp53*$^{-/-}$*Map2k4*$^{mut}$*Cdkn2a*$^{mut}$ background G) Organoid formation. H) Organoid size. n=6 was analyzed per condition, with organoid size measuring n=20 organoids per condition. One-way anova was performed to assess significance. P-value < * = 0.05, <** = 0.01, <*** = 0.001, <****= 0.0001. *Trp53*, abbreviated ("T"), *Nf1* (N), *Map2k4* (M), *Cdkn2a* (C), *Rb1* (R), *Pten* (P), *Brca2* (B2) and *Brca1* (B1).

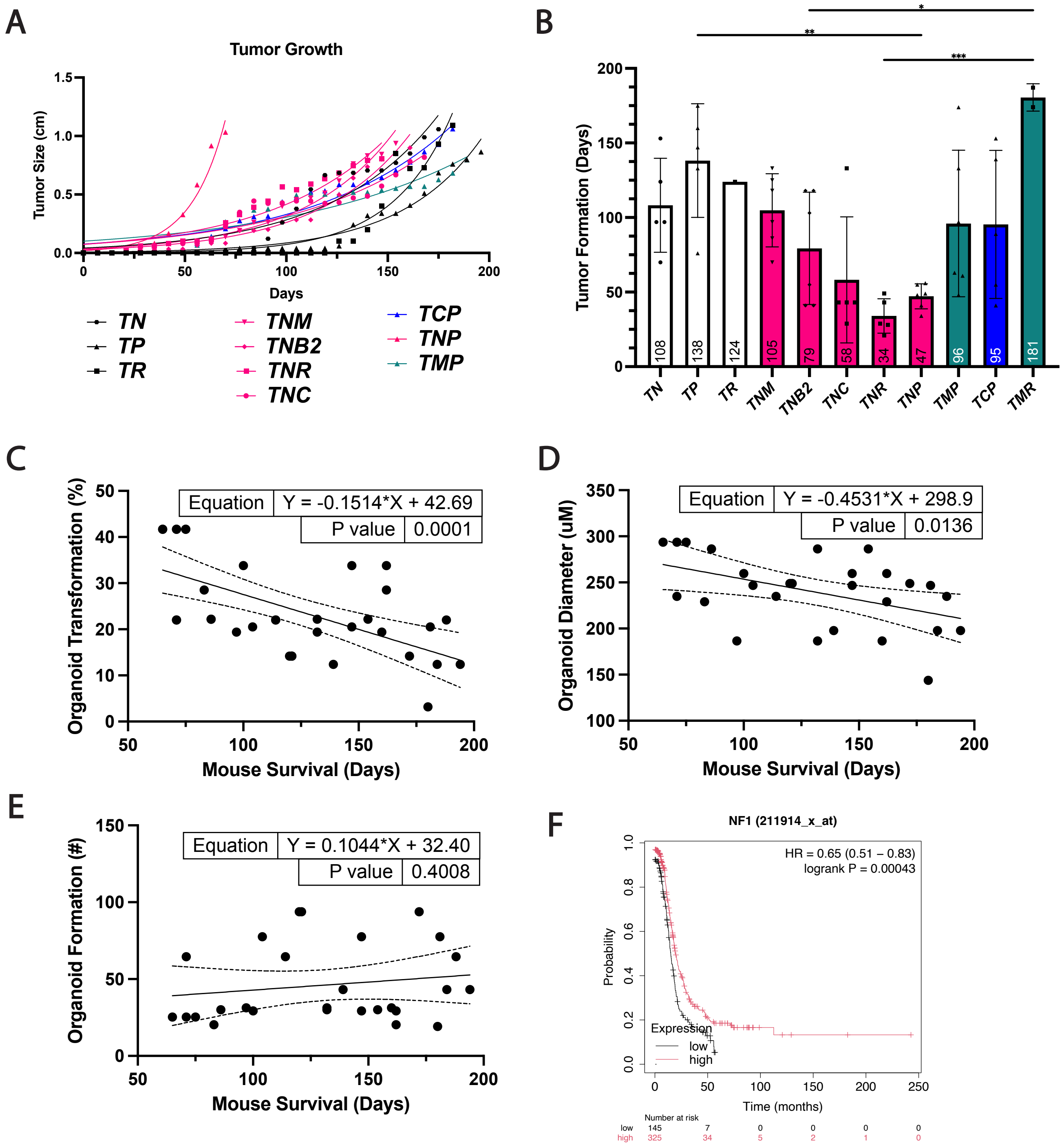


**Supplementary Figure 5. TE organoid transformation reflects tumorigenic potential.** A) Tumor growth as measured on a weekly basis. B) Initial tumor formation, when a palpable mass was first observed. One-way anova was used to assess significance. P-value < * = 0.05, <** = 0.01, <*** = 0.001, <****= 0.0001. C-E) Organoid metrics mapped against mouse survivability. F) Kaplan-Meier survival curve of human patients with high or low *NF1* expression. FDR 5%.

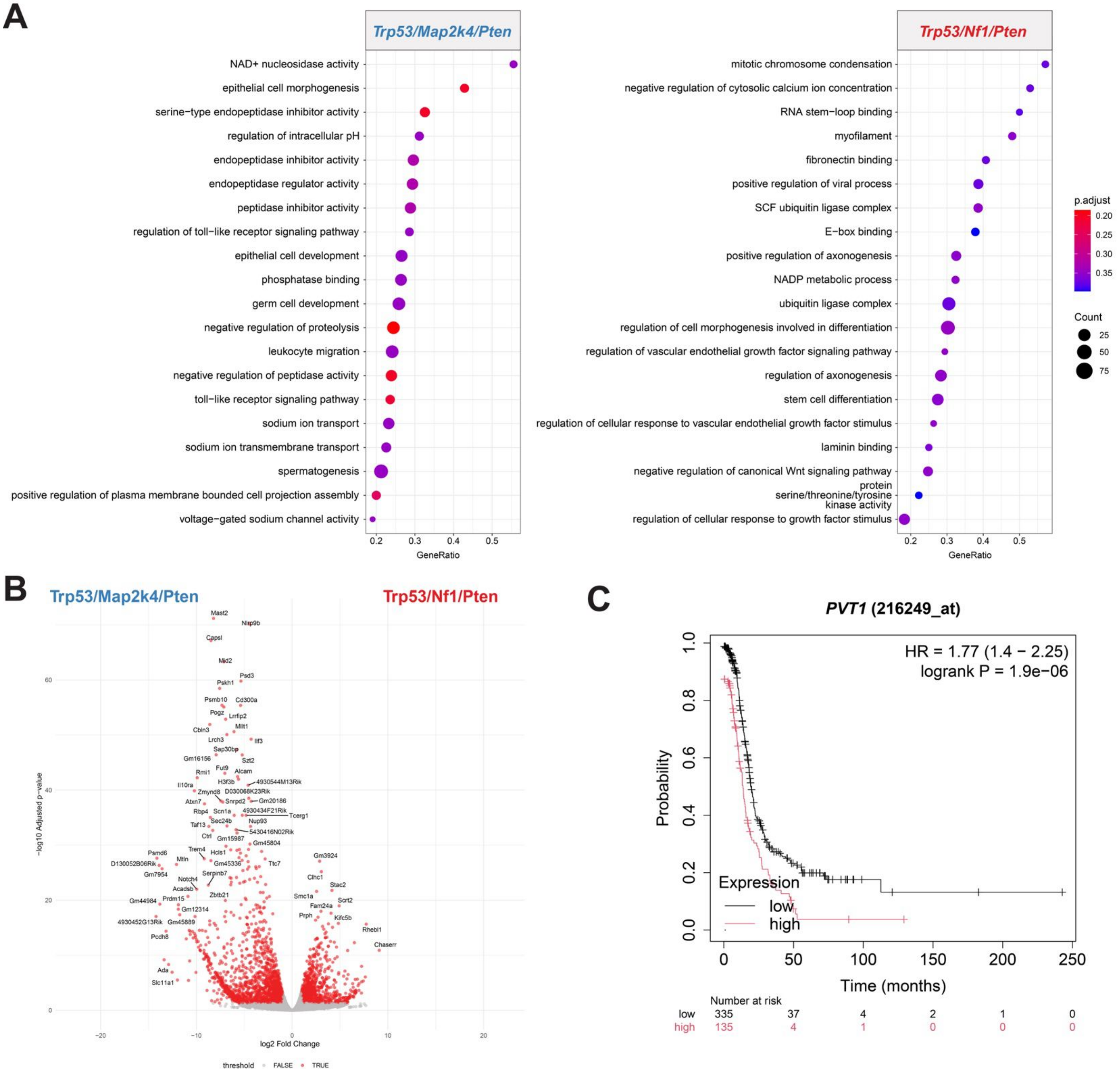


**Supplementary Figure 6. Transcriptomic differences between phenotype and genotype identify *PVT1* as a regulator of survival.** A) GSEA analysis of TMP and TNP. B) Volcano plot of differentially expressed genes between TMP and TNP C) Kaplan-Meier survival curve of human patients with high or low *PVT1* expression. FDR 1%.

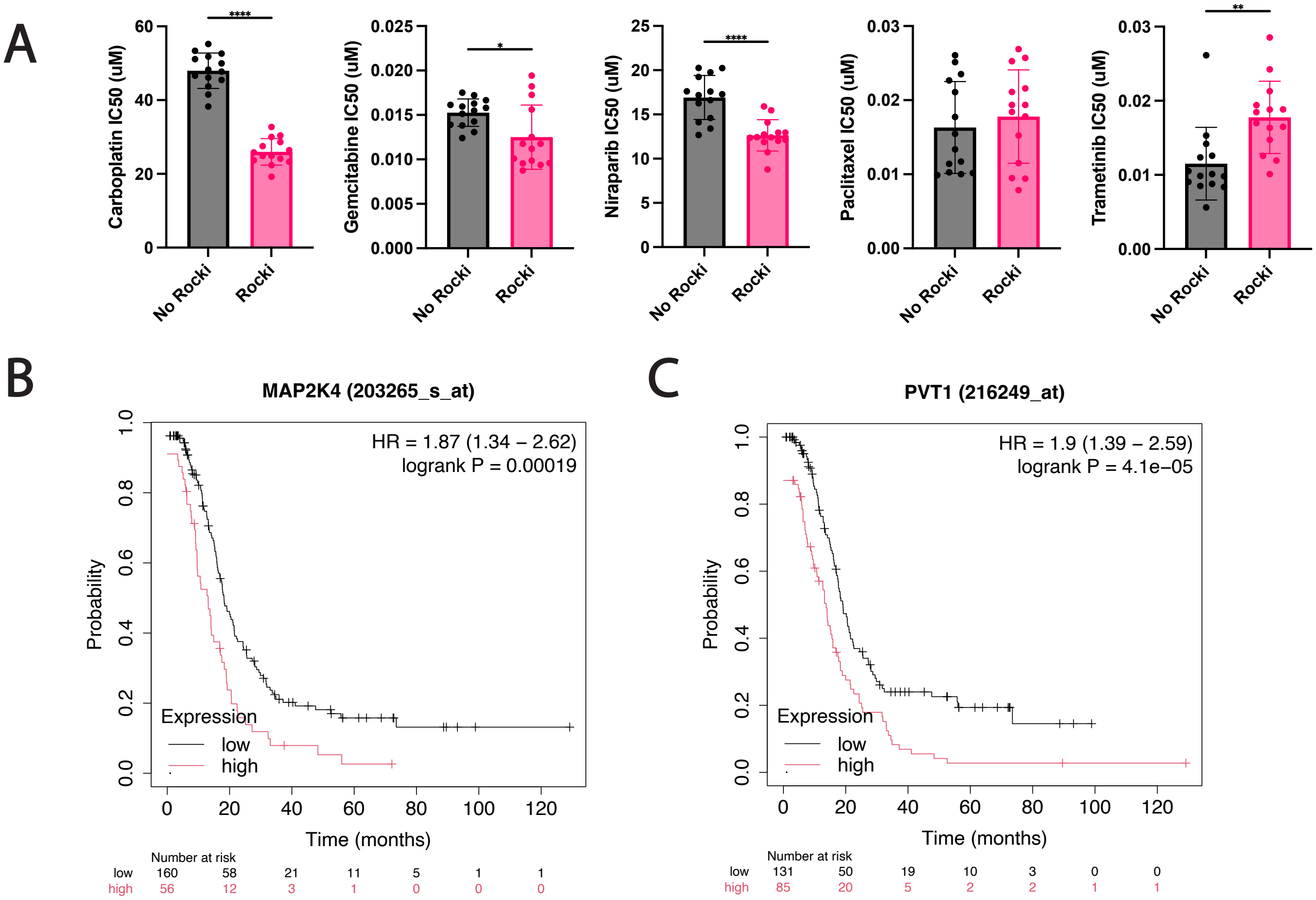


**Supplementary Figure 7. Differences in response to common HGSC therapeutics and human survival.** A) IC50 for all gene combinations with and without ROCKi. Welch's two tailed t-test P-value < * = 0.05, <** = 0.01, <*** = 0.001, <****= 0.0001. Kaplan-Meier survival curve of human patients treated with taxols and improved survival with B) low MAP2K4 expression (FDR 2%) or C) low PVT1 expression (FDR 1%).

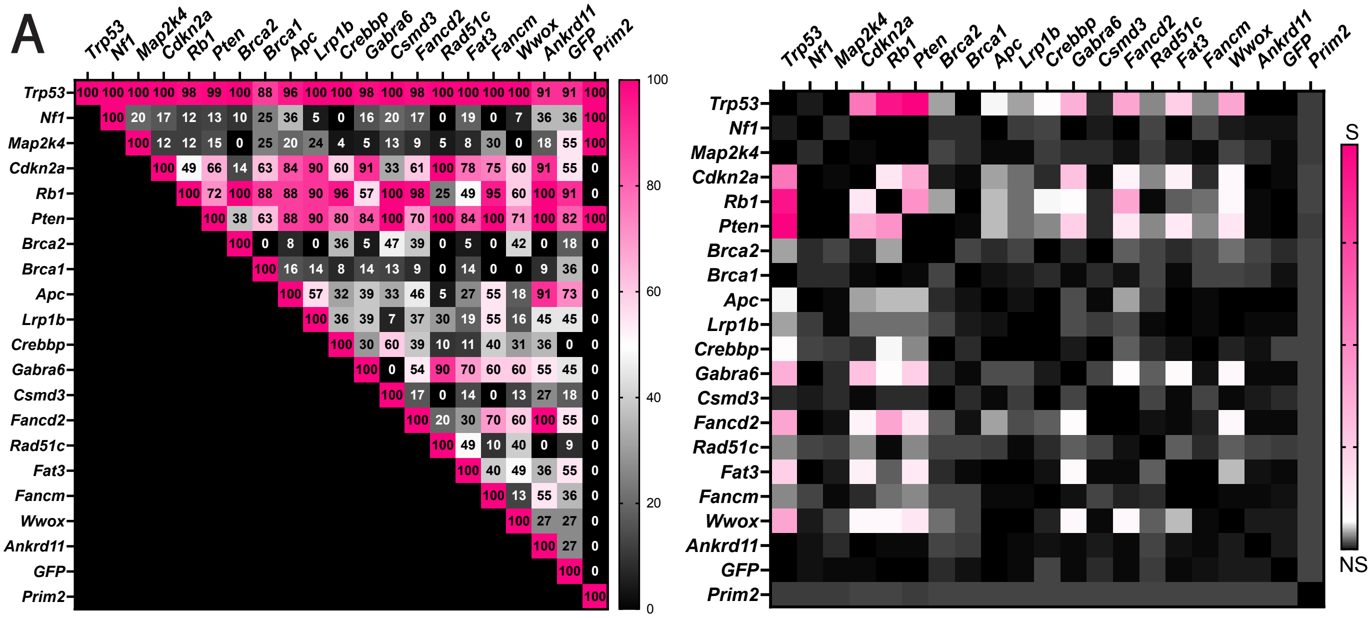


**Supplementary Figure 8**. **Unique and shared drivers of transformation in OSE cells** A) Frequency of co-mutations in OSE colonies, (Right) $\chi^2$ analysis of significant OSE double mutations, DF, 19, P<0.05%. S = Significant, NS = Not Significant.

**Supplementary Table 1 Formation of tumors and their pathology after transplantation of tubal epithelial organoids transformed by combinatorial CRISPR mutagenesis.**

| Targeted genes | Abbreviation | Tumors formed (#) | Mesenchymal | Papillary-glandular | Biphasic |
|---|---|---|---|---|---|
| *Trp53* | T | 0/6 | | | |
| *Trp53/Cdkn2a/Pten* | TCP | 5/6 | | 2 | 3 |
| *Trp53/Map2k4/Brca1* | TMB | 0/6 | | | |
| *Trp53/Map2k4/Brca1* | TMB1 | 1/6 | | 1 | |
| *Trp53/Map2k4/Cdkn2a* | TMC | 0/6 | | | |
| *Trp53/Map2k4/Pten* | TMP | 6/6 | 2 | 4 | |
| *Trp53/Map2k4/Rb1* | TMR | 4/6 | 1 | 3 | |
| *Trp53/Nf1* | TN | 5/6 | 4 | | 1 |
| *Trp53/Nf1/Brca2* | TNB2 | 6/6 | 5 | | 1 |
| *Trp53/Nf1/Cdkn2a* | TNC | 5/6 | 4 | 1 | |
| *Trp53/Nf1/Map2k4* | TNM | 6/6 | 3 | | 3 |
| *Trp53/Nf1/Pten* | TNP | 6/6 | 3 | | 3 |
| *Trp53/Nf1/Rb1* | TNR | 5/6 | | | 5 |
| *Trp53/Pten* | TP | 5/6 | | | 5 |
| *Trp53/Rb1* | TR | 1/6 | | | 1 |

**Supplementary Table 2. Antibodies used for immunostaining.**

| **Antigen, conjugation** | **Antibody source, catalogue number** | **Clone** | **Host** | **Retrieval, Dilution** |
|---|---|---|---|---|
| Acetylated α-Tubulin | Sigma-Aldrich, T7451 | 6-11B-1 | Mouse | Citrate, 1:200 (&IF) |
| FOXJ1 | Novus Biologicals, AF3619-SP | *PC | Goat | Citrate, 1:600 (IF) |
| Ki67 | Invitrogen, 14-5698-82 | SolA15 | Rat | Citrate, 1:4000 (#IHC) |
| Cytokeratin 8, CK8 | Developmental Studies Hybridoma Bank, AB_531826 (20μg $mL^{-1}$) | TROMA-I | Rat | Citrate, 1:50 (IHC) |
| OVGP1 | Abcam, ab118590 | PC | Rabbit | Citrate, 1:600 (IF) |
| P16 | Abcam, ab241543 | PABLO33B | Rat | Citrate, 1:500 (IHC) |
| Pax8 | Proteintech, 10336-1-AP | PC | Rabbit | Citrate, 1:4000 (IHC), 1:200 (IF) |
| Wilm's Tumor Antigen, WT1 | Abcam, ab267377 | EPR23963-116 | Rabbit | Citrate, 1:500 (IHC) |
| tdTomato/RFP | Rockland Immunochemicals Inc., 600-401-379S | PC | Rabbit | No retrieval, 1:4000 (IHC) |
| Anti-rabbit IgG, biotinylated | Vector Labs, BA-1000-1.5 | PC | Goat | 1:200 |
| Anti-rat IgG, biotinylated | Vector Labs, BA-4000-1.5 | PC | Rabbit | 1:200 |

*PC, polyclonal
&IF, immunofluorescence
#IHC, immunoperoxidase-ABC Elite method
